# Cyclase-generated 3’,5’-cGMP activates PucsarTIR antiphage defense through a stepwise zig-zag aligned filament assembly

**DOI:** 10.64898/2026.08.12.744502

**Authors:** Arpita Chakravarti, Yi Wu, A. Maxwell Burroughs, Zhiying Zhang, Shirin Fatma, Raven H. Huang, L. Aravind, Franklin L. Nobrega, Dinshaw J. Patel

## Abstract

Bacteria use nucleotide second messengers to couple phage recognition to antiviral effector activation. Here, we identify Pucsar, an ofshoot of the Pycsar-related antiphage family, whose cyclases generate 3’,5’-cGMP from GTP, that are associated with Toll/Interleukin-1 receptor (TIR) and transmembrane (TM) effectors. In the PucsarTIR system, cGMP produced by cyclase PucC, selectively activates the cognate PucTIR NADase. X-ray crystallography shows that PucC adopts a dimeric cyclase scaffold with a guanine-compatible base-recognition pocket. Cryo-EM and mutational analyses revealed that apo-PucTIR, composed of cyclic nucleotide binding and TIR domains, is self-sequestered as an inactive tetramer, in which TIR domains are proximal but misoriented. Sequential cGMP binding remodels this assembly through partially occupied intermediates and stabilizes a fully cGMP-bound dimer, that polymerizes into a zig-zag aligned filament, creating composite NAD^+^-binding pockets required for efficient NADase activity. These findings establish 3′,5′-cGMP as an antiphage second messenger and reveal how cyclic purine signaling converts self-sequestered TIR effectors into a NADase-competent filament.

## Introduction

Bacteria encode diverse defense systems that detect phage infection and convert this recognition into an immune response^1,2^. One recurring strategy across the tree of Life is the use of nucleotide second messengers, which allows virus sensing by one component to be coupled to the activation of a separate effector module^3–5^. This principle is central to several antiphage systems, including CBASS^6^, Thoeris^7^, bacterial cGAS-STING-like pathways^6^, and CARF-associated CRISPR-Cas systems^8–10^, which use nucleotide-derived signals to trigger antiviral activities. Because many of the activated effectors are potentially toxic to the host cell, these pathways must also tightly regulate when and where effector activity is released.

In addition to these systems, the cyclic mononucleotides were also predicted to function as second messenger signals in bacterial antiphage systems^11^.This was confirmed with the recent characterization of the Pycsar (Pyrimidine cyclase system for antiphage resistance) systems^12^. In these systems, Pycsar cyclases synthesize 3’,5’-cUMP and 3’,5’-cCMP, which activate distinct downstream effectors including CNBD-TIR (cyclic nucleotide binding domain-Toll/Interleukin-1 receptor; PycTIR) and the pJV1-spdB3 family three transmembrane (3TM; PycTM) proteins^12^. PycTIR activation leads to NAD^+^ depletion, whereas PycTM has been associated with membrane-associated antiphage activity^12^. Structural and biochemical studies of PycTIR showed that cUMP is recognized by the N-terminal CNBD and promotes oligomerization required for TIR-dependent NADase activation^13^. However, because the TIR domain was not present in the reported cUMP-bound PycTIR-CNBD structure, how cyclic mononucleotide binding is coupled to full-length CNBD-TIR remodeling, filament assembly, and NADase active-site formation has remained incompletely understood.

These findings established cyclic pyrimidine mononucleotides as antiviral signals^12,13^, but left open the question of whether related phage-defense systems can use cyclic purine mononucleotide counterparts as dedicated antiphage messengers. Cyclic purine mononucleotides, including cAMP and cGMP, are among the best studied second messengers in physiological contexts, regulating processes such as transcription, metabolism, ion-channel activity, and signaling pathways across the tree of Life^14,15^. Despite their broad physiological ubiquity, their role as dedicated antiphage signals has remained unclear.

Comparative genomic studies have identified adenylyl/guanylyl cyclase-like proteins in predicted biological conflict systems, including operons in which cyclase-like enzymes are linked to several distinct predicted and known effectors, with either catalytic or TM domains^16^. These observations suggested the existence of a broader class of cyclic mononucleotide-dependent defense systems, but the identity of the signals produced by these enzymes, the range of associated effectors, and their mechanisms of activation remained unresolved.

To investigate this possibility, we systematically analyzed bacterial cNMP cyclase family proteins from predicted antiphage defense loci. This revealed a diverse set of systems in which the cyclases are recurrently linked to multiple distinct predicted effectors, including CNBD-TIR proteins and pJV1-spdB3 family 3TM proteins. Among these, we identified a branch of cNMP cyclases that we refer to as Pucsar, for purine cyclase system for antiphage resistance. Comparable to the Pycsar systems^12^, these cyclases are commonly linked to CNBD-TIR (PucTIR) and pJV1-spdB3 family 3TM effectors (PucTM)^2^.

We then investigated representative PucsarTIR and PucsarTM systems. We show that cyclases associated with both types of systems generate 3’,5’-cGMP from GTP, establishing Pucsar as a cGMP-producing branch of Pycsar-related antiphage systems. In the PucsarTIR branch, the *Pseudomonas fluorescens* cyclase *Pf*PucC produces 3’,5’-cGMP, which selectively activates the cognate CNBD-TIR effector *Pf*PucTIR to hydrolyze NAD^+^. X-ray structural analysis of *Pf*PucC shows a conserved dimeric cyclase scaffold compatible with GTP recognition, while cryo-EM structures of full-length *Pf*PucTIR capture successive states linking second messenger recognition to NADase activation. In the absence of cGMP, the *Pf*PucTIR dimer is self-sequestered in an inactive dimer-of-dimers tetrameric alignment in which TIR domains are proximal but misoriented. cGMP binding remodels this assembly initially through partially occupied intermediate-like (1:2 and 2:4 cGMP:PucTIR) states and subsequently stabilizes a fully cGMP-bound dimer (2:2 cGMP:PucTIR) that adopts a zig-zag filament in which stacking of adjacent TIR domains creates the composite NAD^+^-binding pockets required for NADase activity. In parallel, a PucsarTM system provides antiphage defense and its associated cyclase produces cGMP, indicating that cGMP-producing cyclases can also signal membrane-associated effectors. The PucTM adopts a cylindrical hexameric fold in the apo state. Together, these findings establish 3’,5’-cGMP as an antiphage second messenger and define how cGMP sensing can be translated into TIR NADase activation through regulated higher-order filament assembly.

## Results

### Comparative genomics and phylogenetic analysis reveal a vast distribution of antiphage cNMP cyclases, including a novel Pycsar-related branch

While the Pycsar systems established that dedicated bacterial antiphage effectors can be activated by cCMP or cUMP^12^, the complete range of nucleotides utilized by related systems remains unclear^2^. Hence, we selected cNMP cyclases from diverse known and predicted anti-selfish element conflict (immune) systems^11^ and their relatives identified by iterative sequence searches for a systematic analysis of their phylogenetic relationships and genomic contexts. To discriminate those that might play immune roles, we also included representative versions of cNMP cyclases that play physiological roles and lack associations with biological conflict effectors (**Table S1**). When we overlaid the conserved gene-neighborhood contexts on the resulting phylogenetic tree, we found that those with potential immune-related versions formed two distinct clades (immunity clades 1 and 2), each with its own distinct outgroup of physiological cNMP cyclases (**Figure 1A**). We then tested the mean branch lengths from the root to the tips of these two predicted immunity-related clades and found that they were significantly longer than those of their respective outgroups (**Figure 1B**, p < 10^-16^). This suggested that they were rapidly evolving relative to their respective outgroups – a feature indicative of accelerated evolution due to being in potential arms races with invasive selfish elements (e.g., viruses)^17^. This observation, taken together with their domain architectures and gene neighborhoods (**Figure 1A**), strongly indicated that these two clades indeed play a role in immune functions.

**Figure 1.**
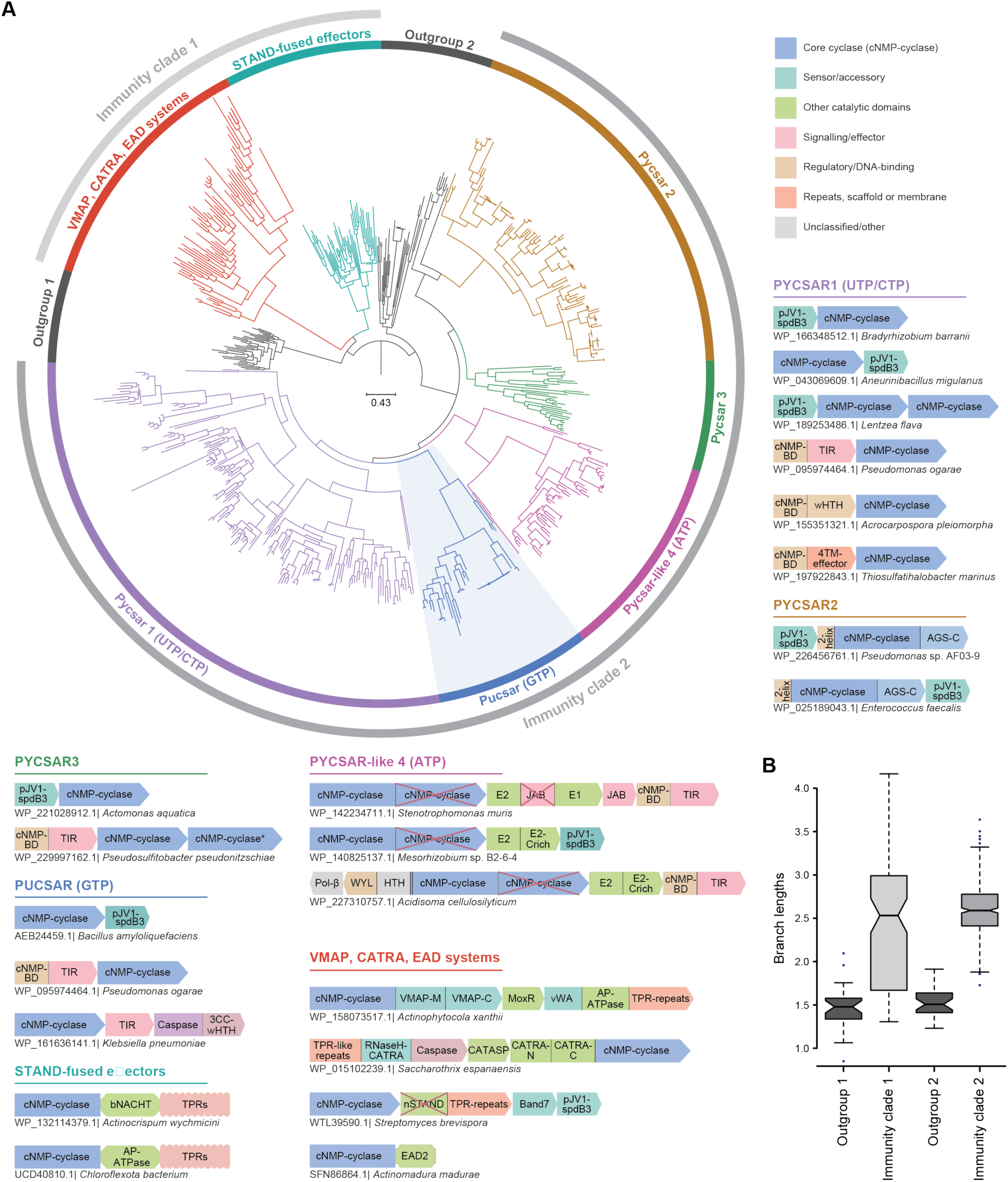
Evolutionary relationships between bacterial cNMP-cyclases involved in immune signaling. **(A)** Phylogenetic tree of the cNMP cyclase domain. Monophyletic clades are colored and labeled with names and known nucleotide substrates, where applicable. Two broader immunity-associated clades are indicated by outer grey arcs. Each of the colored clades has a FastTree clade support value > 90%. The scale bar indicates the average number of substitutions per site. Representative domain architectures of cNMP-cyclases with known or predicted roles in biological conflict are shown adjacent to the clades in which they occur. Individual domains are labeled and colored according to broad functional categories, as indicated in the key. Domain architectures of cNMP cyclase domain-containing proteins are depicted with their individual domains represented by distinct shapes. The cNMP cyclase domain is colored blue for consistency across all architectures. Inactivated enzymatic domains are superimposed with a red ‘X’. Abbreviations: cNMPBD, cNMP-binding domain; TPR, tetratricopeptide repeat; VMAP, vWA-MoxR Associated Protein; CATRA, Caspase and TPR repeat-associated; EAD, Effector-associated domain; TIR, Toll-like immunity receptor; Pol-β, Polymerase-β-nucleotidyltransferase superfamily domain; wHTH, winged helix-turn-helix. **(B)** Boxplots comparing the branch lengths of immune-related cNMP-cyclases with those of their respective outgroups. In each case, the mean branch lengths of the immune-related cNMP-cyclases are significantly greater than those of their outgroups (p < 10^-16^). The accessions corresponding to the tips of the tree are available in **Table S1**.

One of these clades (Immune clade 1 in **Figure 1A**) showed domain fusions with STAND NTPases, such as NACHT and AP-ATPase, a hallmark feature of immune systems across the tree of Life^18^, as well as associations with several less-understood predicted immune systems^11^. The second clade (Immune clade 2 in **Figure 1A**) included the Pycsar system cNMP cyclases and their as yet uncharacterized relatives. This clade further split into five major subclades, each with distinct domain architectures and/or gene-neighborhood associations (**Figure 1A**). Across these clades, two effector classes dominated: cNMP-binding TIR proteins and the 3TM pJV1-spdB3 family.

Of the 5 major subclades, the large Pycsar-1 subclade was composed mainly of standalone cNMP cyclases, most often associated with pJV1-spdB3 family 3TM effectors and less frequently with CNBD-TIR proteins (**Figure 1A**). Some Pycsar-1 loci encoded multiple cyclases in the same genomic region, raising the possibility that these systems may use relay-like or locally amplified cNMP signaling. Two further subclades, Pycsar-2 and Pycsar-3, showed a strong bias in associating with the transmembrane effector, with Pycsar-2 showing this association to the exclusion of the CNBD-TIR effector. The Pycsar-2 subclade cNMP cyclases are further defined by fusions to an N-terminal bihelical domain and a C-terminal AGS-C (Adenylyl/Guanylyl and SMODS C-terminal sensor) domain (**Figures 1A, B**).

Pycsar-like-4 subclade was distinguished by its genomic context, specifically its association with genes encoding E2 ubiquitin-ligase-like proteins^19^. These E2-like proteins are often fused to an inactive JAB domain and sometimes to E1-like domains. They might also be further associated with a gene coding for a JAB peptidase, a domain previously associated with ubiquitin-like protein processing and activation (**Figures 1A, B**)^19^. Some Pycsar-like-4 loci were further linked to regulatory modules containing WYL domains or WYL-Polymerase β (WYL-Polβ) gene pairs. WYL domains are nucleotide-responsive regulatory domains commonly found in prokaryotic defense systems^19^, whereas Polβ-like domains are polymerases that might generate an additional regulatory nucleotide in these systems. These associations suggest that the regulation of Pycsar-like-4 systems might have involved ligation to target proteins mediated by the E1-E2 ligases.

Interestingly, we found that a fifth uncharacterized subclade, hereinafter the Pucsar subclade, formed a distinct sister group to the Pycsar-1 subclade (**Figure 1A**). Pucsar loci showed associations with both major effector classes, namely CNBD-TIR proteins and pJV1-spdB3-like 3TM effectors of antiphage defense (**Figures 1A, B**). Some PucsarTIR loci also encoded additional predicted effectors, including caspase superfamily peptidase, whereas rare loci contained alternative effector candidates, such as luciferase-like monooxygenases, enzymes associated with oxidative chemistry^20^. Thus, Pucsar systems represent a distinct branch of Pycsar-related antiphage cyclases. We next asked whether the phylogenetic and architectural diversification of these cyclases were accompanied by differences in predicted nucleotide specificity. Mapping subclade-specific residues onto structural models further revealed distinct residue patterns around the nucleotide-binding pocket, consistent with diversification of substrate preference across the major Pycsar-related cyclase lineages (**Figure S1A**). We generated AlphaFold3^21^ models of representative cyclases from each subclade with alternative nucleotide substrates and divalent ions and compared the atom-by-atom plDDT scores of the modelled ligands (**Table S2**). Most Pucsar cyclases showed higher scores for GTP than UTP, supporting a predicted preference for a purine substrate. In contrast, Pycsar-1, Pycsar-2, and Pycsar-3 cyclases predominantly favored UTP over GTP, consistent with the experimentally established UTP preference of characterized Pycsar uridylate cyclases^12,13^. Pycsar-like-4 cyclases showed a distinct preference for ATP (**Table S2**). Together, these analyses suggest diversification of nucleotide specificity across the major Pycsar-related cyclase subclades, and in particular, predict a preference for GTP among Pucsar cyclasess

### Representative Pucsar systems provide antiphage defense through 3’,5’-cGMP signaling

Having identified Pucsar as a distinct branch of Pycsar-related cyclase systems, we next asked whether representative Pucsar loci function as antiphage defense systems in cells. We selected systems corresponding to the two major effector branches identified by comparative genomics: a *Pseudomonas fluorescens* PucsarTIR locus encoding a cyclase, here referred to as *Pf*PucC, and a cognate CNBD-TIR effector, here referred to as *Pf*PucTIR (**Figure 2A**, top panel); and an *Escherichia coli* PucsarTM locus encoding a cyclase (*Ec*PucC) associated with a pJV1-spdB3-like TM3 effector (*Ec*PucTM), (**Figure 2B**, top panel). Both Pucsar systems showed antiphage activity, although with different spectra and strengths. The PucsarTIR system reduced infection by siphophage c1a (**Figure 2A**, bottom panel), whereas the PucsarTM system showed broader activity against multiple phages, particularly T4-like phages (**Figure 2B**, bottom panel).

**Figure 2.**
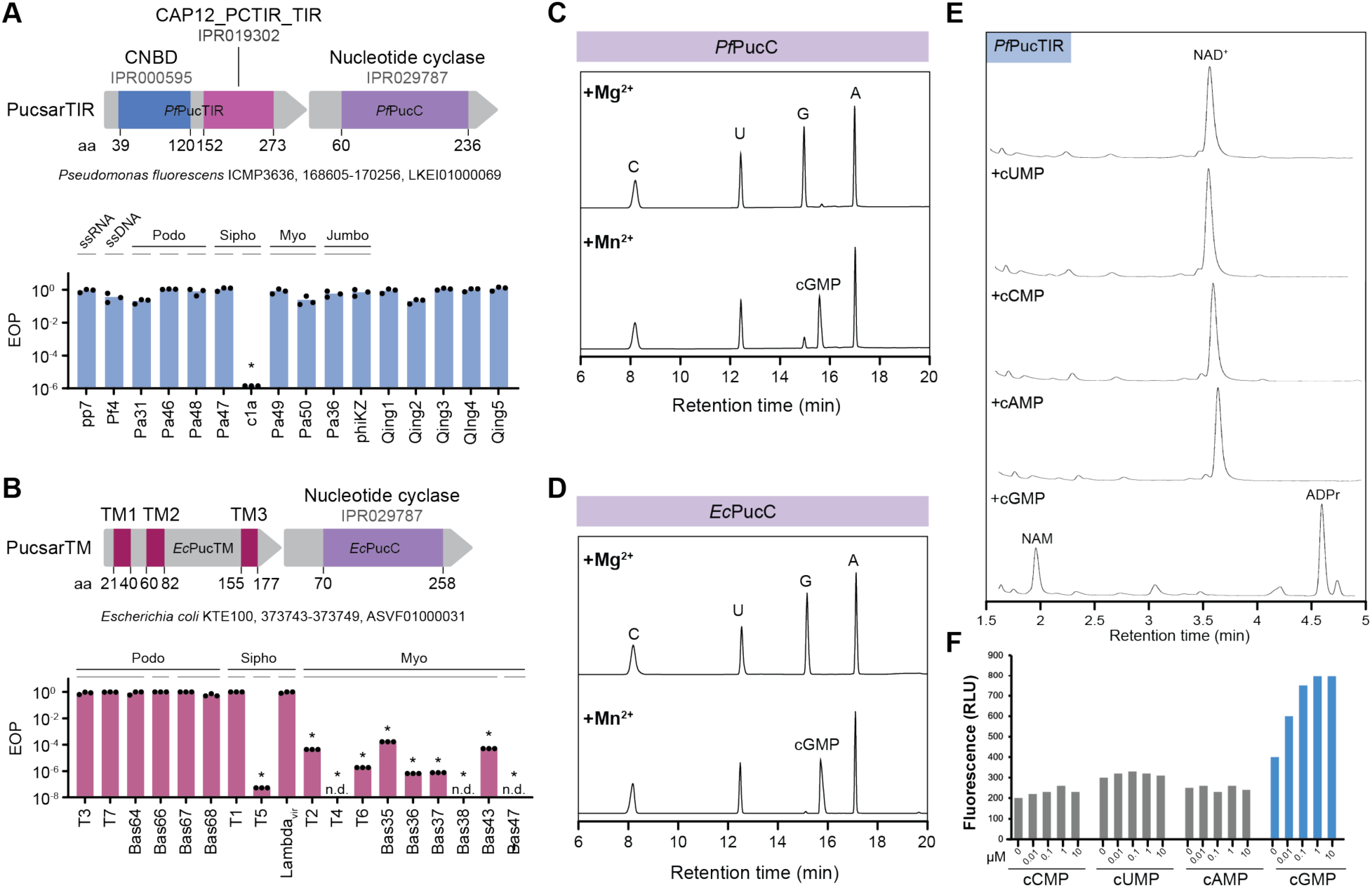
Representative Pucsar systems provide antiphage defense and generate 3’,5’-cGMP. **(A)** Domain organization of the *Pseudomonas fluorescens* PucsarTIR locus, endocing the CNBD-TIR effector *Pf*PucTIR and the nucleotide cyclase *Pf*PucC (top panel), and efficiency of plating (EOP) assays measuring antiphage activity across a panel of phages in *Pseudomonas aeruginosa* strain PAO1 (bottom panel). **(B)** Domain organization of the *Escherichia coli* PucsarTM locus, encoding the transmembrane effector *Ec*PucTM and the nucleotide cyclase *Ec*PucC (top panel), and EOP assays measuring antiphage activity across a panel of phages in *E. coli* BL21. n.d., no detectable plaques (bottom panel). **(C)** Representative chromatograms of nucleotide products generated by purified *Pf*PucC after incubation with ATP, GTP, CTP, and UTP in the presence of Mg^2+^ or Mn^2+^. **(D)** Representative chromatograms of nucleotide products generated by purified *Ec*PucC after incubation with ATP, GTP, CTP, and UTP in the presence of Mg^2+^ or Mn^2+^. **(E)** Representative chromatograms of NAD^+^ hydrolysis reactions containing purified *Pf*PucTIR and the indicated cyclic nucleotide. **(F)** Fluorescence-based NAD+ depletion assay measuring activation of *Pf*PucTIR by increasing concentrations of cUMP, cCMP, cAMP, or cGMP.

We next sought to identify the cyclic nucleotide signal produced by the Pucsar cyclases. Previous studies of nucleotide-based defense systems have shown that purified cyclases can be reconstituted *in vitro* with nucleotide triphosphate substrates and divalent cations to identify their cyclic nucleotide products^12,13,22,23^. Hence, we recombinantly expressed and purified *Pf*PucC from PucsarTIR and *Ec*PucC from PucsarTM (**Figure S1B**) and incubated each protein with ATP, GTP, CTP, and UTP. Reactions were performed in the presence of Mn^2+^, which supports the *in vitro* activity of Pycsar-related cyclases^12^, with Mg^2+^ included as a control (**Figures 2C, D**). Both cyclases converted GTP into a product with the retention time expected for 3’,5’-cGMP in the Mn^2+^ reaction, with no comparable product under the Mg^2+^ control condition. Co-injection with synthetic 3’,5’-cGMP confirmed that the cyclase-derived product corresponded to 3’,5’-cGMP (henceforth labeld cGMP) (**Figure S1C**), establishing that representative PucsarTIR and PucsarTM cyclases generate cGMP from GTP. This shared product experimentally supports the predicted preference of Pucsar cyclases for GTP and distinguishes the Pucsar cyclases tested here from characterized Pycsar enzymes that generate cyclic pyrimidine mononucleotides^12^.

The specific production of cGMP by *Pf*PucC suggested that this cyclic nucleotide could activate the cognate TIR NADase effector of the PucsarTIR system. To test this, we first measured TIR-dependent NAD^+^ depletion during infection with a PucsarTIR-sensitive phage. Cells expressing the PucsarTIR system showed reduced intracellular NAD^+^ levels compared with infected control cells (**Figure S1D**). Consistent with this *in vivo* phenotype, purified *Pf*PucTIR effector (**Figure S1E**) was selectively activated by 3’,5’-cGMP *in vitro* (**Figure 2E**). Addition of cGMP stimulated NAD^+^ hydrolysis by *Pf*PucTIR, producing ADP-ribose (ADPr) and nicotinamide (NAM) (**Figure S1F**), whereas cAMP, cCMP, and cUMP showed substantially weaker or no activation under the same conditions (**Figure 2E**). Fluorescence-based NAD^+^ depletion assays across a range of cyclic nucleotide concentrations confirmed that cGMP was the most potent activator tested (**Figure 2F**).

Together, these data show that representative PucsarTIR and PucsarTM systems provide antiphage defense and that their associated cyclases generate cGMP. For the TIR branch, cGMP directly links the cyclase to effector activation by stimulating *Pf*PucTIR-dependent NAD^+^ hydrolysis.

### The *Pf*PucC cyclase forms a two-fold symmetric domain-swapped homodimer with interfacial GTP binding pockets

To understand how the PucC cyclase recognizes GTP and catalyzes cyclic nucleotide formation, we purified apo-PucC. The protein eluted as a dimer by size-exclusion chromatography (**Figure S1B**), consistent with the dimeric architecture observed for related adenylyl/guanylyl cyclase-like enzymes and Pycsar cyclases^12^. We determined the crystal structure of apo-PucC at 1.9 Å resolution, revealing a twofold symmetric, domain-swapped homodimer (**Figure 3A**, crystallography data collection and statistics in **Table S3**). Each protomer contributed to a deep pocket at the dimer interface, positioning the predicted active site between the two subunits. Thus, PucC adopts the conserved dimeric cyclase fold used by Pycsar and adenylyl/guanylyl cyclases, despite producing cGMP rather than the cyclic pyrimidine mononucleotides generated by characterized Pycsar enzymes.

**Figure 3.**
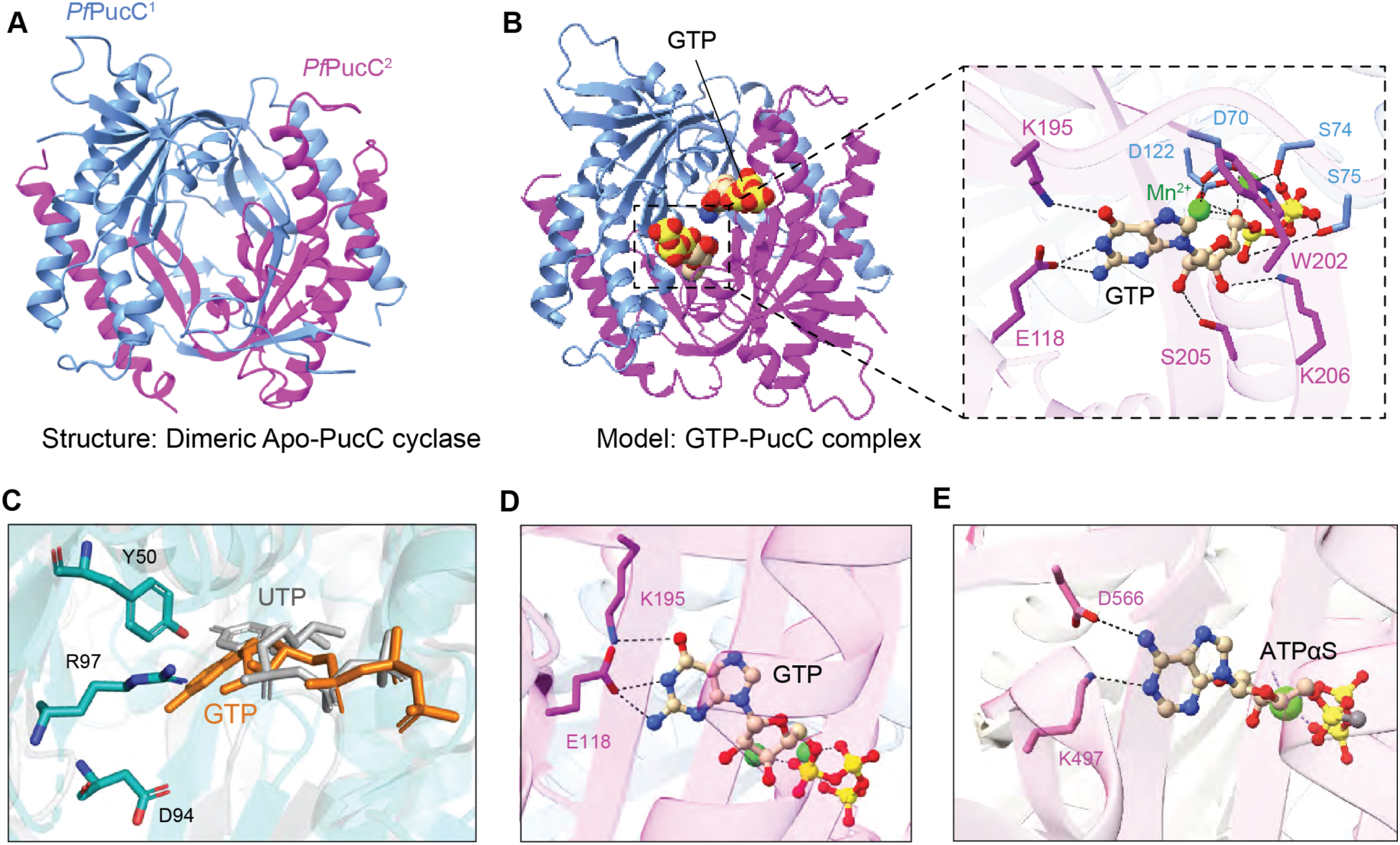
Structural analysis of apo-and GTP-bound *Pf*PucC. **(A)** X-ray crystal structure of apo-*Pf*PucC determined at 1.9 Å resolution, showing a domain-swapped homodimer with twofold symmetry. The two protomers are shown in blue and magenta. **(B)** AlphaFold 3 (AF3) model of the 2:2:4 *Pf*PucC:GTP:Mn^2+^ complex. The model contains two *Pf*PucC protomers, two GTP molecules, and four Mn^2+^ ions, corresponding to two equivalent interfacial active sites. The insert shows the predicted GTP-binding pocket and the residues positioned to contact the nucleotide and coordinate the catalytic metals. **(C)** Superposition of the GTP-binding pocket (in orange) in the AF3 model of 2:2:4 *Pf*cNC:GTP:Mn^2+^ with the UTP-binding pocket (in silver) in the Proteinix model of the 2:2:4 *Bc*PycC:UTP:Mn^2+^, with special emphasis on Arg97, Tyr50 and Asp94 of *Bc*PycC sterically occluding the guanine base. **(D, E)** Comparison of nucleobase recognition in *Pf*PucC (panel D), and *Ca*RhGC E497K/C566D (panel E). In *Pf*PucC, Gly118 and Lys195 form predicted contacts with the guanine base, whereas Lys497 and Asp566 contact the adenine base in *Ca*RhGC E497K/C566D. Metal ions are shown as green spheres, and the predicted polar and metal-coordination interactions as dashed lines.

Because attempts to capture a substrate-bound crystal structure were unsuccessful, we used AlphaFold 3 (AF3)^21^ to model a 2:2:4 PucC:GTP:Mn^2+^ complex, comprising two PucC protomers, two GTP molecules, and four Mn^2+^ ions. This stoichiometry placed one GTP and two Mn^2+^ ions in each interfacial active site (pTM = 0.90, ipTM = 0.90) (**Figure 3B**). To distinguish conserved catalytic features from determinants of substrate specificity, we compared this model with Proteinix^24^ models of 2:2:4 *Bc*PycC:UTP:Mn^2+^ (pTM = 0.96, ipTM = 0.95,) (**Figure S2A**) and 2:2:4 *Ec*PycC:CTP:Mn^2+^ (pTM = 0.89, ipTM = 0.92) (**Figure S2B**)^12^, as well as with the crystal structure of an ATPαS-bound engineered rhodopsin adenylyl cyclase. (**Figure S2C**)^25^.

Across these models and structures, conserved acidic residues contributed to phosphate and divalent cation coordination (**Figure 3B** insert, **Figures S2D-F**), suggesting a shared mechanism for 3’,5’-cyclic mononucleotide synthesis. In the AF3 model of the *Pf*PucC:GTP:Mn^2+^complex, Asp70 and Asp122 were positioned near the phosphate groups and metal ions (**Figures S2G, H**). One Mn^2+^ ion coordinated the β and γ phosphates, whereas the second Mn^2+^ ion coordinated the α phosphate and the ribose 3’-OH-aligned. This arrangement positions the 3’-OH for in-line nucleophilic attack on the α-phosphate, enabling formation of a 3’,5’-cyclic mononucleotide and release of pyrophosphate (schematic in **Figure S2H**). Thus, purine-and pyrimidine-producing members of this cyclase family appear to share a conserved catalytic mechanism for cyclic mononucleotide synthesis.

In contrast, the predicted base-recognition interactions differed markedly between *Pf*PucC and pyrimidine-producing Pycsar cyclases. In the *Bc*PycC:UTP:Mn^2+^ model, residues such as Tyr50, Asp94, and Arg97 formed a restrictive binding pocket that accommodates the smaller pyrimidine base, but would sterically disfavor binding of the larger guanine base (**Figure 3C**). By contrast, the *Pf*PucC:GTP:Mn^2+^ model showed a more permissive interfacial pocket, with Glu118 and Lys195 positioned near and recognizing the Watson-Crick edge of the guanine base (**Figure 3D**). These differences provide a structural rationale for why *Pf*PucC selectively uses GTP and produces cGMP rather than the cyclic pyrimidine mononucleotides generated by characterized Pycsar cyclases.

To further interpret the predicted guanine-recognition site, we compared the Watson-Crick edge recognition positions (E118 and K195) of *Pf*PucC (**Figure 3D**), with K497 and D566 for the adenine-recognition site in the ATPαS-bound structure of an engineered rhodopsin adenylyl cyclase domain (**Figure 3E**)^25^. In the latter structure, residues at the corresponding base-recognition positions create an adenine-binding environment that favors ATP. This comparison supports the idea that substrate specificity in *Pf*PucC is shaped by a small number of base-contacting residues within the interfacial active site, whereas the comparison with Pycsar cyclases explains the broader shift from pyrimidine to purine cyclic mononucleotide production.

Together, these structural and modeling analyses show that *Pf*PucC combines a conserved dimeric cyclase scaffold with a distinct interfacial base-recognition pocket compatible with GTP selection. This provides a structural basis for the biochemical finding that *Pf*PucC generates cGMP and supports the broader prediction that changes in nucleobase-recognition residues underlie signal diversification across Pycsar-related antiphage cyclases.

### Apo-*Pf*PucTIR is self-sequestered in an inactive tetrameric state

Having shown that cGMP activates PucTIR-dependent NAD^+^ hydrolysis, we next asked how this CNBD-TIR effector is maintained in an inactive state before signal binding. We purified apo-PucTIR (**Figure S1E)** and determined a 2.78 Å cryo-EM structure (**Figure 4A**, cryo-EM workflow in **Figure S3A;** cryo-EM statistics listed in **Table S4**). The apo-PucTIR structure revealed a twofold symmetric dimer-of-dimers tetrameric architeture. The assembly contained two internally directed, extended monomers (in magenta and gold) and two externally directed, fold-back monomers (in blue and green) (**Figure 4A**). Comparison of the extended (in magenta) and fold-back (in blue) monomers showed that these represent distinct conformations of the PucTIR effector, with an rmsd of 2.48 Å after alignment of their TIR domains (**Figure 4B**). The tetramer (**Figures 4A** and **S3B**) was stabilized by extensive hydrogen bonding interactions between both the TIR (**Figure S3C**) and CNBD (**Figure S3D**) domains. TIR-TIR contacts connected the internal and external monomers through interactions involving the αT1-αT3 and αT2-αT4 helices, as well as adjacent loop regions (boxed regions in **Figure S3C**). CNBD-CNBD contacts further stabilized the overall dimer-of-dimers architecture, primarily through interactions between paired α-helices αC1-αC2 and αC3-αC4 (boxed regions in **Figure S3D).**

**Figure 4.**
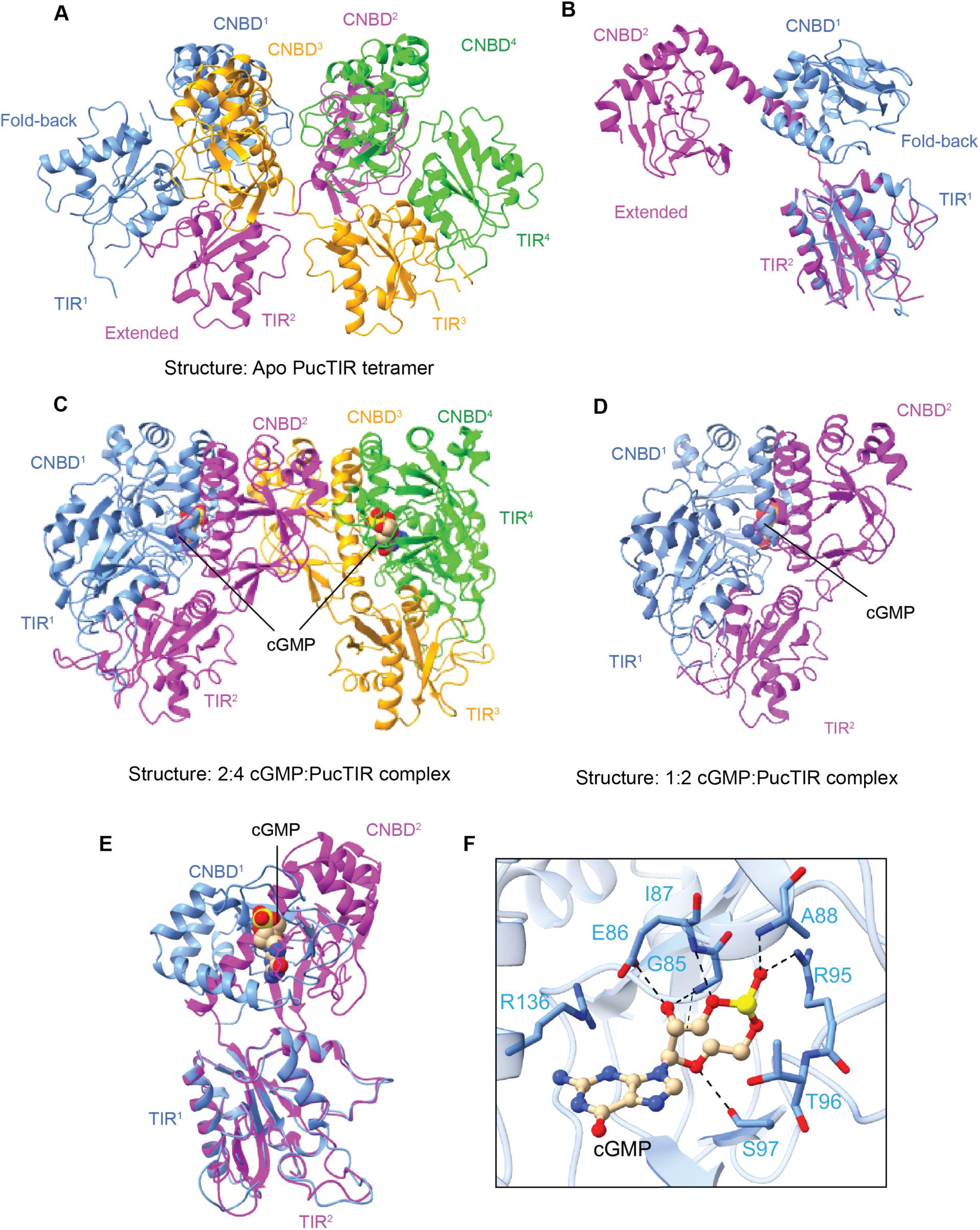
Apo-*Pf*PucTIR forms a self-sequestered tetramer that is remodelled by partial cGMP binding. **(A)** Cryo-EM structure of apo-*Pf*PucTIR determined at 2.78 Å resolution, showing a dimer-of-dimers tetramer. The assembly contains two fold-back protomers (in blue and green) and two extended protomers (in purple and gold), with CNBD and TIR domains indicated for each chain. **(B)** Superposition of an extended (in magenta) and a fold-back (in blue) PucTIR protomer from the apo tetramer, aligned on the TIR domain. The two conformations differ substantially, with an rmsd of 2.48 Å. **(C)** Cryo-EM structure of the partially occupied 2:4 cGMP:PucTIR tetramer determined at 2.8 Å resolution. Two cGMP molecules are bound within the CNBD pockets of the externally oriented fold-back protomers (in blue and green). **(D)** Cryo-EM structure of the partially occupied 1:2 cGMP:PucTIR dimer determined at 3.14 Å resolution. One cGMP molecule is visible in the CNBD pocket of the fold-back protomer (in blue). **(E)** Superposition of an extended (in magenta) and a fold-back (in blue) PucTIR protomer from the 1:2 cGMP:PucTIR complex, aligned on the TIR domain. The two conformations exhibit an rmsd of 2.33 Å. **(F)** Close-up view of the cGMP-binding pocket in the 1:2 cGMP:PucTIR dimer. Residues positioned around the guanine base, ribose, and cyclic phosphate are shown as sticks, with cGMP shown in ball-and-stick representation.

Importantly, apo-*Pf*PucTIR was catalytically inactive in the absence of cGMP. Although the TIR domains are positioned close to one another in the apo tetramer, their relative orientation does not generate an NADase-competent composite active site (**Figure 4A**). Thus, the apo *Pf*PucTIR represents a preassembled but self-sequestered state, in which the effector domains are held in a constrained arrangement that prevents productive NAD^+^ binding and hydrolysis. This mode of inhibition is distinct from most other TIR domain containing immune proteins, whereby the inactivity is often achieved by keeping the TIR domains spatially separated^26^, prohibiting reconstitution of active NADase pockets.

### Partial cGMP binding remodels the *Pf*PucTIR tetramer but does not form an active NADase site

Having established that apo-PucTIR dimer is held in a self-sequestered tetrameric state, we next asked how cGMP binding begins to remodel this inactive assembly. We therefore incubated PucTIR with cGMP and analyzed the complex by cryo-EM under conditions that yielded two cGMP-bound reconstructions: a 2.8 Å 2:4 cGMP:PucTIR tetramer containing two cGMP molecules bound across four PucTIR protomers **(Figure 4C)**, and a 3.14 Å 1:2 cGMP:PucTIR dimer containing one cGMP molecule bound across two protomers **(Figure 4D**), with the cryo-EM work flow shown in **Figure S4A** and cryo-EM statistics listed in **Table S4** for both complexes. The partially occupied 1:2 cGMP:PucTIR dimer (**Figure 4D**) resembled one half of the partially occupied 2:4 cGMP:PucTIR tetramer (**Figure 4C**), with cGMP bound selectively to the fold-back but not extended protomer in both complexes. Because these structures contain cGMP in only one of the two available CNBDs, they appear to capture partially occupied states along the activation pathway, rather than the fully active assembly. Superposition of the cGMP-bound fold-back (in blue) and unbound extended (in magenta) protomers in these partially occupied cGMP-PucTIR complexes yielded an rmsd = 2.33 Å after alignment of their TIR domains (**Figure 4E**).

The bound cGMP was positioned in the CNBD pocket through specific contacts with the guanine ribose sugar, and cyclic phosphate backbone (**Figure 4F**). Arg136 was positioned against the guanine base, consistent with a cation-π interaction that may help stabilize ligand binding, whereas the cyclic phosphate was anchored by multiple polar contacts. However, we did not observe any guanine base-specific contacts in this partially occupied state. This suggests that the intermediate captures an early cGMP-bound state in which ligand binding has begun to remodel the tetramer, but before the fully occupied, activated architecture is stabilized.

Notably, while the cGMP-bound external fold-back protomer exhibited an rmsd of 1.4 Å when comparing the apo-and partially cGMP-bound *Pf*PucTIR structures after alignment of the TIR domains (**Figure S4B**), comparison of the two internally oriented extended protomers between apo-and partially cGMP-bound *Pf*PucTIR structures exhibited a much larger rmsd of 2.32 Å (**Figure S4C**). Thus, cGMP binding to the fold-back pair of protomers (**Figure S4B**) is associated with long-range remodeling of the extended pair of protomers (**Figure S4C**), indicating that ligand binding is transmitted allosterically through the assembly, rather than acting only locally within the occupied CNBDs.

The partially occupied 1:2 cGMP:PucTIR dimer (**Figure 4D**) resembled one half of the partially occupied 2:4 cGMP:PucTIR tetramer (**Figure 4C**) and contained a single cGMP molecule bound to one (fold-back) of the two CNBDs. This suggests that partial cGMP occupancy can promote not just remodeling, but also dissociation, of the apo-like dimer-of-dimers tetrameric architecture into smaller cGMP-bound dimeric states. Although the 1:2 cGMP:PucTIR dimer sample was prepared in the presence of the non-cleavable NAD^+^ analog BAD (benzamide adenine dinucleotide), we did not observe clear density for BAD or NAD^+^-derived products in either reconstruction. Together with the partial cGMP occupancy, this indicates that early cGMP-bound conformations do not form a stable substrate-bound, NADase-competent state under these conditions. These structures therefore define intermediate-like states in which cGMP has begun to remodel the self-sequestered tetramer, including dissociation to a dimeric state, but full effector activation has not yet occurred.

### Full cGMP occupancy assembles *Pf*PucTIR into a filament built from cGMP-bound dimers

The partially occupied structures suggested that cGMP binding can remodel the apo-tetramer, but that early ligand-bound states are not sufficient to form an NADase-competent complex. We therefore collected cryo-EM data and analyzed an additional *Pf*PucTIR sample incubated with cGMP under modified buffer conditions (250 mM NaCl compared with 150 mM used previously) to capture further cGMP-bound conformations. In this dataset, cryo-EM micrographs and classification revealed filamentous *Pf*PucTIR particles (**Figure 5A**). A 3.34 Å cryo-EM reconstruction of these particles showed that the filament is built from repeating two-fold symmetric *Pf*PucTIR dimers (**Figure 5B**), with each protomer of the dimer containing a cGMP in its CNBD pocket (cryo-EM work flow in **Figure S5A** and cryo-EM statistics in **Table S4**).

**Figure 5.**
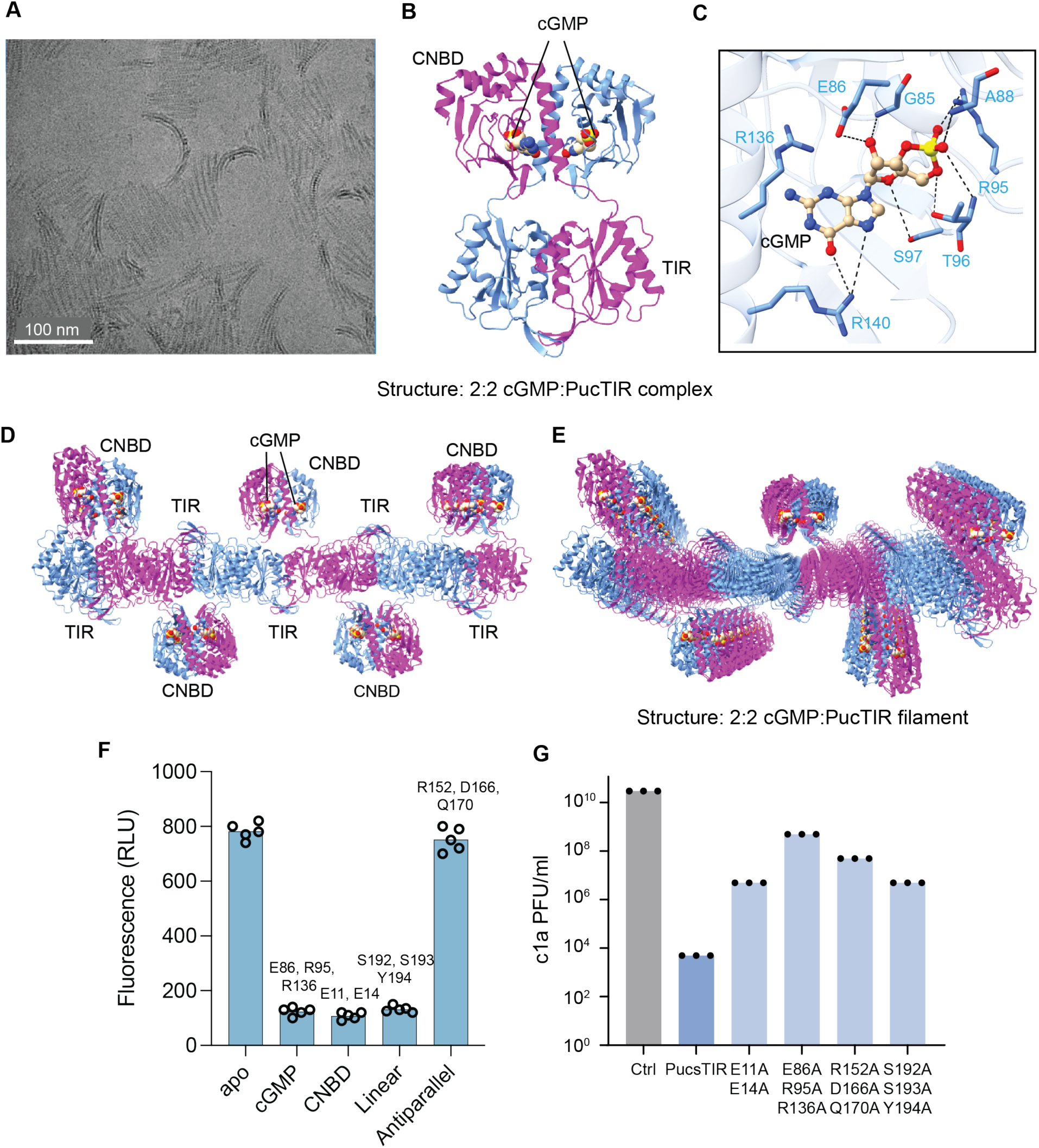
Full cGMP occupancy assembles *Pf*PucTIR into a filament required for antiphage activity. **(A)** Representative cryo-EM micrograph showing curved PucTIR filaments formed in the presence of cGMP. Scale bar, 100 nm. **(B)** The cryo-EM structure of the fully occupied 2:2 cGMP:PucTIR dimer, the basic repeating unit of the linear filament, determined at 3.34 Å resolution. One cGMP molecule is bound to the CNBD of each protomer. **(C)** Close-up view of the cGMP-binding pocket in the fully occupied 2:2 cGMP:PucTIR. Residues positioned around the guanine base, ribose and cyclic phosphate are shown as sticks, with cGMP shown in ball-and-stick representation. **(D)** A single layer of zig-zag aligned CNBD and TIR domains of fully occupied 2:2 cGMP:PucTIR dimer toplogy. The TIR domains interact with each other along a central spine, while the CNBD domains are aligned in opposing directions, reflective of a zig-zag alignment. **(E)** A view of the filament formed by the 2:2 cGMP:PucTIR dimer, following stacking of indiviual dimer units in the context of the zig-zag filament alignment. **(F)** *In vitro* fluorescence-based NAD⁺ depletion assay measuring cGMP-stimulated NADase activity of wild-type PucTIR and structure-guided mutants targeting the cGMP-binding pocket, CNBD-CNBD interface, linear filament contacts, or antiparallel zig-zag interface. The data are shown for five replicates. **(G)** *In vivo* antiphage activity of PucsarTIR variants carrying the indicated PucTIR mutations, measured as phage production after infection. Mutations affecting cGMP binding, dimer stabilization, linear filament contacts, or antiparallel zig-zag packing reduce antiphage defense *in vivo*. The data are shown for three replicates.

The repeating unit of the filament was a twofold symmetric PucTIR dimer containing two cGMP molecules, with one cGMP bound to each CNBD (**Figure 5B**). This 2:2 cGMP:*Pf*PucTIR complex therefore represents a fully occupied dimeric repeat, in contrast to the partially occupied 1:2 cGMP:PucTIR complex (**Figure 4D**) captured in the previous dataset. Thus, the two cGMP-bound datasets appear to capture different states along the same activation pathway: an intermediate state in which only one of two CNBDs is visibly occupied (**Figure 4D**), and a filament-forming state in which both CNBDs within each dimeric repeat are occupied (**Figure 5B**).

The cGMP-binding mode in the filament-forming dimer (**Figure 5C**) was similar but not identical to that observed for the fold-back protomer in the partially occupied cGMP-bound structures (**Figure 4F**). As in the earlier complexes, Arg136 stacked over the guanine base, while the ribose and cyclic phosphate were anchored by a network of polar interactions. However, the fully occupied 2:2 cGMP:PucTIR complex also revealed an additional interaction in which Arg140 contacted the O6 and N7 atoms along the Hoogsteen edge of the guanine base (**Figure 5C**). This interaction was not observed in the intermediate 1:2 or 2:4 cGMP:PucTIR structures (**Figure 4F**), suggesting that full cGMP occupancy stabilizes a more defined ligand-recognition state.

Full cGMP occupancy was accompanied by a major rearrangement of the CNBD-TIR architecture. In the filament-forming 2:2 cGMP:PucTIR complex, both symmetry-related *Pf*PucTIR protomers adopted an extended, domain-swapped conformation, with interactions between paired CNBD domains and paired TIR domains (**Figure 5B**). Superposition of the TIR domains showed that this extended conformation differed substantially from the cGMP-bound fold-back protomer (rmsd = 5.6 Å; **Figure S5B**), and from the unbound extended protomer (rmsd of 5.7 Å; **Figure S5C**) observed in the partially occupied 1:2 cGMP-bound dimer. Thus, binding of cGMP to both CNBDs is associated with conversion of the effector from a partially remodeled state into an extended dimeric conformation, that is compatible with filament assembly (**Figures 5D,E**).

### Adoption of a zig-zag alignment by the 2:2 cGMP:*Pf*PucTIR on higher-order filament formation

Cryo-EM micrographs of the 2:2 cGMP:PucTIR complex exhibited curved and helical filament-like architecture (**Figure 5A**). Processing and refinement of the 3.34 Å cryo-EM data set of the complex (**Figure S5A**) yielded a zig-zag aligned filament topology (**Figures 5D, E**). This topology was composed of linear filaments involving stacked basic 2:2 cGMP:PucTIR dimeric units, which were further organized in a zig-zag arrangement mediated by central TIR-TIR domain interactions, with CNBD domains directed in opposite directions. There are extensive interactions between CNBD domains and between TIR domains not only within individual dimers (**Figure 5B**), but also between stacked dimers in the context of the linear filament, as well as between TIR domains within adjacent linear filaments (**Figure 5E**), thereby forming a super-higher order zig-zag alignment.

Together, these structures support a stepwise model for PucTIR activation. Initial cGMP binding remodels the sequestered apo dimer-of-dimers tetrameric architecture (**Figure 4A**), facilitating dissociation into a dimer (**Figure 4D**), but does not generate a fully assembled effector architecture. Full cGMP occupancy then stabilizes an extended domain-swapped dimer (**Figure 5B**) that polymerizes into a zig-zag aligned filament (**Figure 5E**). This defined sequence of conformational transitions provides a structural basis for how the cGMP signal produced by the PucC cyclase is coupled to higher-order assembly of the PucTIR effector.

### Linear *Pf*PucTIR filament contacts are required for efficient NADase activity

Having established that the 2:2 cGMP:PucTIR complex forms a higher-order zig-zag aligned filament (**Figure 5E**) built from two-fold symmetric dimeric repeats (**Figure 5B**), we next asked which interfaces within this assembly are required for NADase activity. Inspection of the filament structure revealed two levels of higher-order organization. First, the fully occupied 2:2 cGMP:PucTIR dimeric repeats stack to form a linear filament. Second, neighboring linear filaments associate through additional TIR-TIR contacts, producing a zig-zag arrangement (**Figure 5E**). Thus, the filament structure raised the question of whether NADase activity depends specifically on formation of the linear fragment, and whether the higher-order zig-zag packing contributes to the stability of filament formation.

Comparison of the the dimeric topologies within the apo (**Figure 6A**), partially cGMP-bound, (**Figure 6B**) and fully cGMP-bound (**Figure 6C**) structures are supportive of a stepwise model in which cGMP binding remodels PucTIR from an inactive tetramer into a filament-forming dimeric repeat. In this overview, partial cGMP binding first remodels the tetrameric topology therby destabilizing the apo dimer-of-dimers arrangement, but partial occupancy is not sufficient to generate the active architecture. Full occupancy of cGMPs by both CNBDs stabilizes the extended dimeric repeat, allowing these repeats to stack into the linear filament as part of the zig-zag higher order alignment. This suggested that residues involved in cGMP binding, dimer stabilization, and linear filament contacts should all be important for NADase activation.

**Figure 6.**
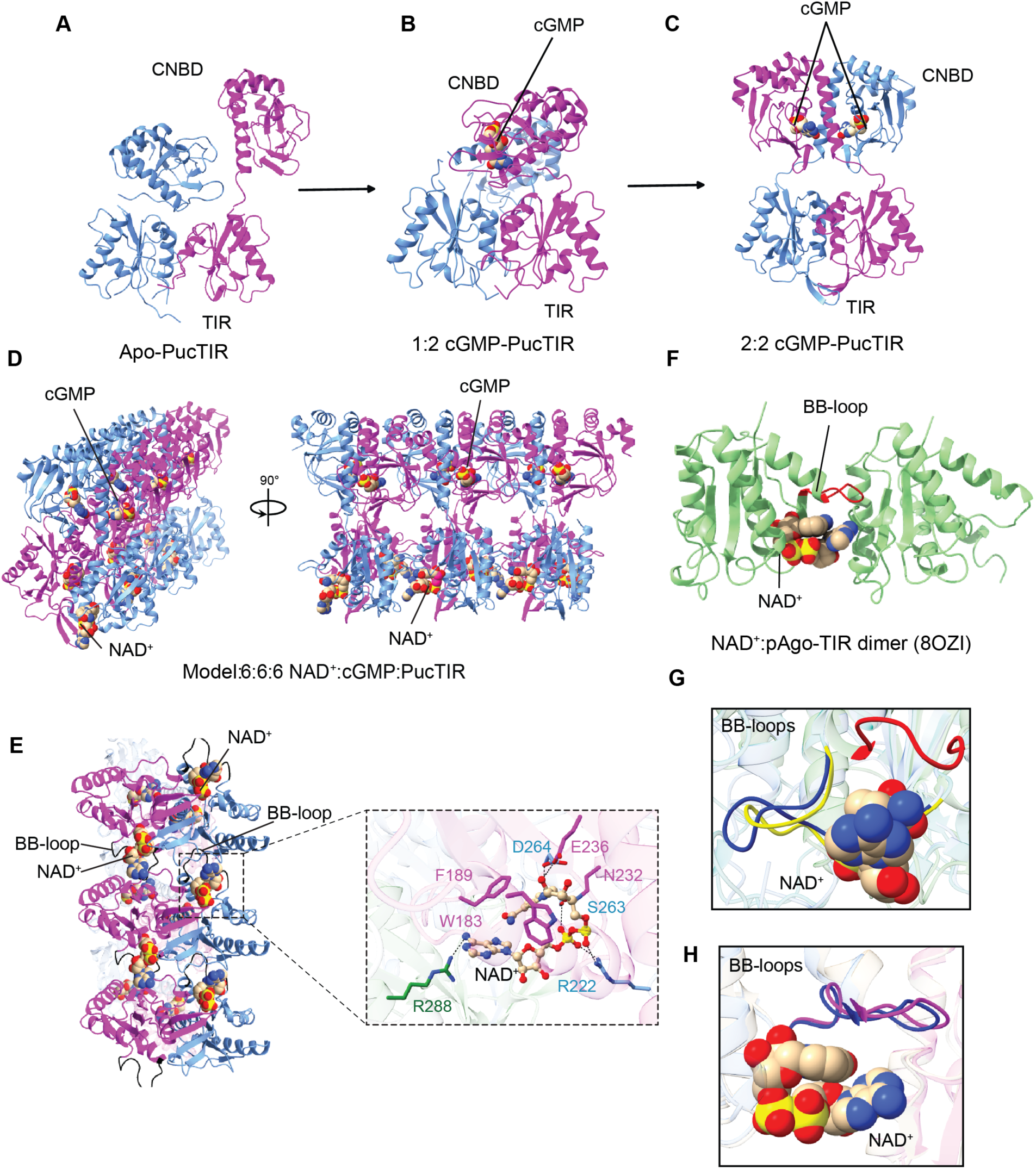
Model for stepwise cGMP-dependent activation of *Pf*PucTIRs NADase activity with linear filament formation predicted to create an accessible composite NAD^+^-binding site in *Pf*PucTIR. (A-C) Dimeric alignments in the self-sequestered inactive apo-*Pf*PucTIR tetramer (panel A). Initial cGMP binding remodels the tetramer and releases a partially occupied dimeric 1:2 cGMP:PucTIR intermediate state (panel B). Full cGMP occupancy stabilizes a 2:2 cGMP-PucTIR dimeric repeat (panel C) that assembles into a linear filament. **(D)** A Proteinix model of a substrate-bound PucTIR linear filament segment containing six PucTIR protomers, six cGMP molecules and six NAD^+^ molecules. The model retained cGMP in the experimentally observed CNBD pockets and placed NAD^+^ at interfaces between TIR domains of adjacent stacked 2:2 cGMP:PucTIR dimers. **(E)** View of the Proteinix model of cGMP-and NAD^+^-bound PucTIR filament, highlighting the predicted positions of NAD^+^ molecules between stacked TIR domains. The insert shows a predicted composite NAD^+^-binding pocket formed by residues contributed by neighboring TIR domains. **(F)** NAD^+^-bound pAgo-TIR dimer structure (PDB: 8OZI), showing the position of the BB loop away from the bound NAD^+^, leaving the substrate-binding site accessible. **(G)** Comparison of BB-loop positions in the NAD^+^-bound pAgo-TIR dimer (in red), apo-*Pf*PucTIR (in blue) and the partially occupied 1:2 cGMP:PucTIR state (in yellow). In apo (in blue) and partially occupied 1:2 cGMP:PucTIR complex (in yellow), the BB loop occupies the region corresponding to access to the NAD^+^-binding pocket, consistent with a nonproductive or occluded substrate-binding state. **(H)** Comparison of BB-loop positions in the experimentally determined 2:2 cGMP:PucTIR linear filament (in blue) and the Proteinix model of the NAD^+^-bound 2:2 cGMP:PucTIR filament (in magenta). In both structures, the BB loop is displaced away from the predicted NAD^+^-binding pocket, consistent with an accessible substrate-binding site.

To test this model, we purified structure-guided *Pf*PucTIR mutants designed to perturb distinct features of the cGMP-bound assembly. These included mutations in the cGMP-binding pocket (E86A/R95A/R136A; **Figures S6A, E**), the CNBD-CNBD interface within the dimeric repeat (E11A/E14A; **Figures S6B, F**), TIR-domain contacts within the linear filament (S192A/S193A/Y194A; **Figures S6C, G**), and the higher-order zig-zag interface between neighboring linear filaments (R152A/D166A/Q170A; **Figures S6D, H**). We then monitored cGMP-stimulated NAD^+^ hydrolysis using a fluorescence-based NAD^+^ depletion *in vitro* assay. Mutations in the cGMP-binding pocket (E86A/R95A/R136A) strongly reduced NAD^+^ hydrolysis (**Figure 5F**), consistent with the requirement for cGMP-dependent activation. Disruption of the CNBD-CNBD interface (E11A/E14A) also reduced activity (**Figure 5F**), indicating that stabilization of the fully occupied dimeric repeat contributes to effector function. Importantly, mutations targeting TIR-domain contacts within the linear filament (S192A/S193A/Y194A) strongly reduced NAD^+^ hydrolysis (**Figure 5F**), showing that stacking of cGMP-bound dimers into the linear filament is required for efficient NADase activity. In contrast, mutations predicted to disrupt the antiparallel zig-zag higher order alignment interface (R152A/D166A/Q170A) did not substantially alter NAD^+^ hydrolysis (**Figure 5F**). Interestingly, however, all mutants reduced the *in vivo* antiphage activity of PucsarTIR, including mutations in the zig-zag interface (**Figure 5G**). This suggests that the zig-zag interface is not required for catalytic activation under purified *in vitro* conditions, but contributes to defense *in vivo*.

These results indicate that the linear *Pf*PucTIR filament possesses the relevant architecture for NADase activation, whereas the zig-zag packing between neighboring linear filaments is dispensable for NAD^+^ hydrolysis *in vitro* but contributes to antiphage defense in cells. This difference suggests that zig-zag packing alignment of the higher-order filament may influence PucTIR function under cellular conditions, for example by affecting filament stability, organization, localization, or productive activation during infection.

### *Pf*PucTIR filament formation is predicted to create an accessible composite NAD^+^-binding site

The requirement for linear filament contacts suggested that filament assembly may create the catalytic architecture needed for PucTIR NADase activity. However, although samples prepared with cGMP and the NAD^+^ analogue BAD formed filament-like particles, we did not observe clear density for BAD, NAD^+^, or NAD^+^-derived products in the cryo-EM maps, which reflected linear filament, but not zig-zag filament formation. Unfortunately, the TIR domains were more flexible in the reconstructions, limiting direct interpretation of substrate binding. This indicated that the substrate-bound state is either transient, heterogeneous, or not stably captured under the conditions tested. We therefore used structure-guided modelling calculations to ask how the cGMP-bound linear PucTIR filament could position NAD^+^ in the compoite pocket formed at TIR-TIR interfaces.

To model a possible substrate-bound state, we generated a linear PucTIR filament segment containing six PucTIR protomers, six cGMP molecules, and six NAD^+^ molecules using Proteinix. The model (ipTM = 0.64, pTM = 0.61) retained cGMP in the experimentally observed CNBD pockets and placed NAD^+^ at composite sites formed between TIR domains from adjacent stacked dimeric repeats (**Figure 6D**).

In the predicted substrate-bound filament, each NAD^+^ molecule was positioned at an interface formed by multiple TIR domains (**Figure 6E** and insert). Arg288 was positioned near the nicotinamide-ribose region of NAD^+^, consistent with a possible role in substrate recognition or stabilization, while neighboring TIR domains contributed additional residues predicted to contact the adenine base and phosphate groups. This arrangement provides a structural rationale for why the linear filament is required for NADase activity: stacking of cGMP-bound dimers creates composite TIR interfaces that are absent in the apo tetramer (**Figure 4A**), the 1:2 cGMP:*Pf*PucTIR complex (**Figure 4D**) or in the isolated dimeric repeat of the 2:2 cGMP:*Pf*PucTIR complex (**Figure 5B**).

We next compared these predicted NAD^+^-binding sites with experimentally characterized TIR NADase structures to assess whether the modeled pocket is compatible with known catalytic conformations. A key structural element in this comparison is the BB loop, a flexible region of the TIR domain, that can regulate access to the NAD^+^-binding pocket. In inactive TIR-domain arrangements, this loop can occlude the substrate-binding site, whereas activated assemblies often reposition the BB loop away from the entrance to the catalytic pocket^27^.

We used NAD^+^-bound pAgo-TIR dimer (PDB: 8OZI) as a reference because it provides a direct view of an accessible NAD^+^-binding site in an active TIR interface. In this structure, the BB loop is positioned away from the NAD^+^-binding pocket (**Figure 6F**, and in red, **Figure 6G**, in red), leaving the substrate site accessible. By contrast, in both the apo-*Pf*PucTIR tetramer (**Figure 6G**, in blue) and the partially 1:2 cGMP-PucTIR intermediate (**Figure 6G**, in yellow), the BB loop projects into the region corresponding to the predicted NAD^+^-binding site, which could sterically occlude substrate access. In the fully-bound 2:2 cGMP-PucTIR filament, however, the BB loop is repositioned away from the predicted NAD^+^-binding site (in magenta, **Figure 6H**), resembling the open configuration observed in the NAD^+^-bound pAgo-TIR dimer (in red, **Figure 6G**). This movement is also preserved in the Proteinix model of the NAD^+^-bound *Pf*PucTIR filament, in which the BB loop does not block entry into the predicted substrate-binding pocket (in blue, **Figure 6H**).

Together with the mutational data, these observations suggest that cGMP-dependent filament formation activates *Pf*PucTIR by creating composite NAD^+^-binding pockets between stacked TIR domains of 2:2 cGMP:PucTIR and by repositioning the BB loop to make these pockets accessible. This model explains why mutations that disrupt linear filament contacts, impairs NAD^+^ hydrolysis, while mutations affecting higher-order zig-zag filament packing do not.

### The PucsarTM effector forms a cylindrical hexameric membrane assembly

Having defined the mechanism of the PucsarTIR branch, we next examined the structurally distinct transmembrane effector associated with Pucsar. As shown above, the *E. coli* PucsarTM system provides broad antiphage protection, particularly against T4-like phages (**Figure 2B**, lower panel), and its associated cyclase *Ec*PucC generates cGMP from GTP (**Figure 2D**). To investigate the architecture of the membrane effector, we purified *Ec*PucTM composed of CNBD and TM domains in LMNG/GDN (lauryl-maltose neopentyl glycol/glyco-diosgenin) detergent (**Figure S7A**) and determined its apo cryo-EM structure at 2.67 Å resolution (cryo-EM workflow in **Figures S7B, C**).

*Ec*PucTM assembled into a six-fold symmetric cylindrical complex with distinct membrane-spanning and cytosolic regions (**Figure 7A-C**). Each protomer contains three transmembrane helices (labeled TM1, TM2 and TM3 in magenta) (**Figure 2B**, top panel) together with a predominantly α-helical cytosolic region (in blue) and a β-sheet-containing appendage (also in blue). The six TM2 α-helices are directed toward the centre of the assembly (**Figure 7B**), defining a central membrane-spanning channel approximately 15.2 Å in diameter (**Figure 7C**). Six unassigned densities are located towards the inside of the hexameric pore likely corresponding to detergents. This density lacked defining features for a specific chemical orientation, and therefore, was excluded from the final atomic model.

**Figure 7.**
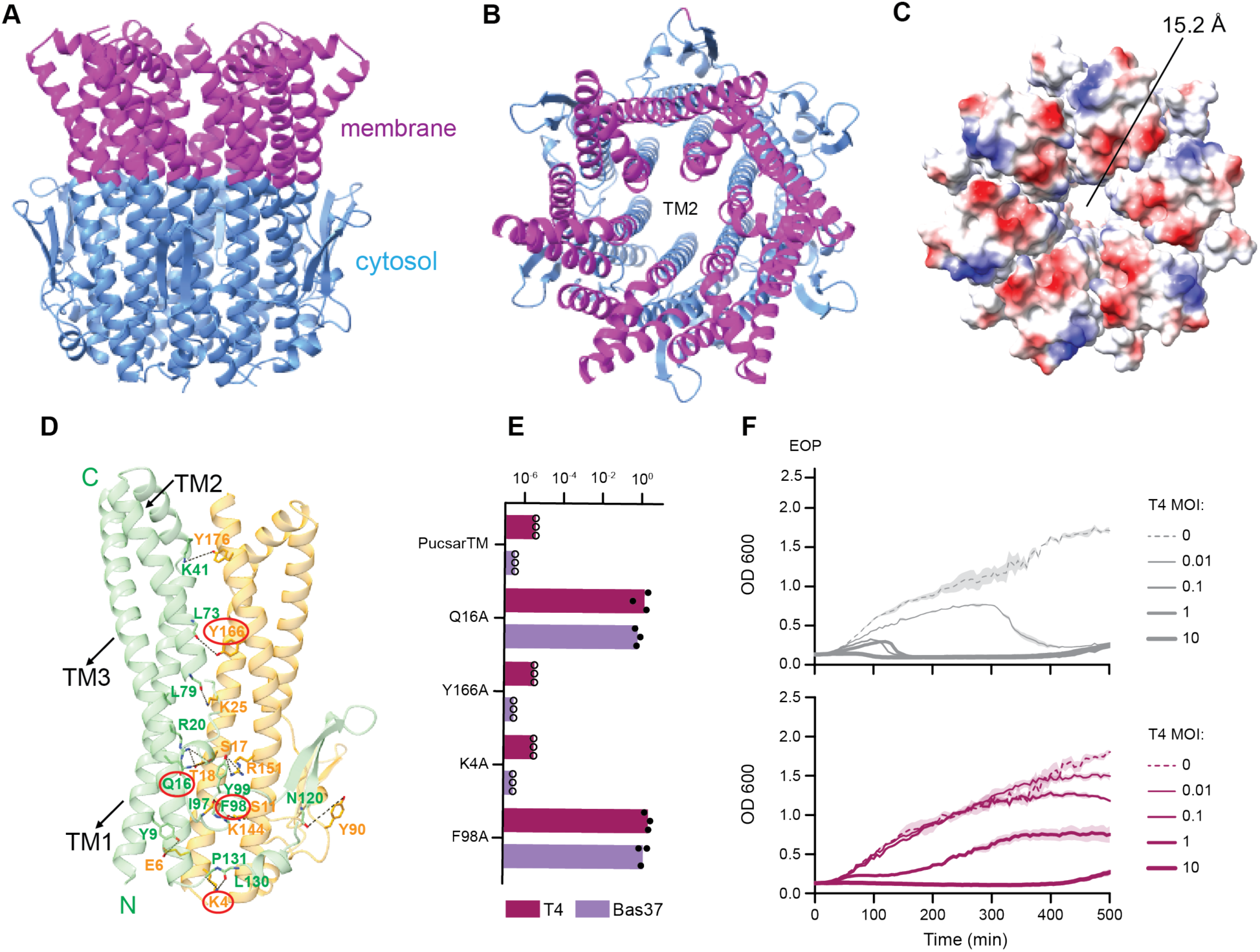
*Ec*PucTM forms a hexameric membrane assembly required for antiphage defense. **(A, B)** Side and membrane-facing views of the 2.67 Å cryo-EM structure of apo-*Ec*PucTM, which assembles into a six-fold symmetric cylindrical complex. Membrane-spanning and cytosolic regions are shown in magenta and blue, respectively. The TM2 α-helices are directed toward the centre of the hexamer and line the central membrane-spanning channel as shown in panel B. **(C)** Electrostatic surface representation of *Ec*PucTM viewed from the membrane side, showing the central channel, with an approximate diameter of 15.2 Å. **(D)** Close-up view of the interface between two adjacent *Ec*PucTM protomers, highlighting intersubunit contacts within the transmembrane and cytosolic regions. Residues selected for structure-guided mutagenesis are encircled. **(E)** Efficiency of plating (EOP) of the indicated phages on cells expressing wild-type PucsarTM or PucsarTM carrying the indicated *Ec*PucTM alanine substitutions. Individual points represent biological replicates. **(F)** Growth of cells expressing PucsarTM following infection with phage T4 at low and high multiplicities of infection (MOI). Uninfected cultures were analyzed in parallel.

Extensive contacts between neighbouring protomers are observed across both the transmembrane and cytosolic regions of the cylindrical hexameric assembly (**Figure 7D**). To examine the functional importance of residues contributing to these intersubunit interfaces, we selected residues positioned at or near interfaces between adjacent *Ec*PucTM protomers for structure-guided mutagenesis. Alanine substitutions were then tested for their effects on antiphage activity. Substituion of Q16 or F98 strongly impaired PucsarTM-mediated defense against the tested phages, whereas Y166A and K4A retained activity close to that of the wild-type system (**Figure 7E**). Thus, only a subset of the tested interfacial residues were required for defense, identifying specific positions within the *Ec*PucTM assembly that are important for its antiviral function.

Our attempts to generate and solve the cryo-EM structure of cGMP bound to PucTIR have not been successful to date and will require further experimentartion to be brought to fruition. We have undertaken Proteinix-based modelling of the cGMP-PucTIR complex but are uncertain about the validity of positioning of the cGMP within the cyclindrical hexameric PucTM scaffold in the computed structures at this time.

Finally, we monitored bacterial growth following T4 phage infection across a range of multiplicities of infection. PucsarTM-expressing cultures showed an MOI-dependent response to infection, with gro’wth strongly impaired at high MOI but maintained or recovered at lower MOIs (**Figure 7F**). This infection-dependent phenotype was consistent with an abortive infection response, in which protection of the bacterial population is associated with growth arrest of heavily infected cells.

Together, these findings show that the Pucsar family couples cGMP-producing cyclases to structurally distinct antiviral effectors: a filament-forming TIR NADase in the PucsarTIR family and a hexameric membrane-associated effector in the PucsarTM family.

## Discussion

Characterized Pycsar systems use cyclic pyrimidine mononucleotides, such as cCMP and cUMP, to regulate the deployment of antiviral effectors^12,13^. By contrast, the representative Pucsar systems described here feature cyclases that generate cGMP. This establishes cGMP as an antiphage second messenger and suggests that diversification of the cNMP cyclase active site has allowed Pycsar-related antiphage systems to explore distinct cyclic nucleotide signals. The observation that Pucsar cyclases are linked to both PucTIR (CNBD-TIR) and PucTM (3TM) effectors further suggests that cGMP-producing cyclases can be coupled to multiple antiviral outputs. Within the Pucsar subclade, our investigation of the PucsarTIR effectors provides a direct structure-based mechanistic link between cGMP production and effector activation.The *Pf*PucC cyclase preserves the domain-swapped homodimeric fold and catalytic framework observed in Pycsar cyclases and other classical physiological cNMP cyclase-enzymes^12,13,25^. In particular, conserved acidic residues are positioned to coordinate divalent cations that, in turn, interact with the phosphate groups and are required for 3’,5’-cyclic mononucleotide synthesis. However, the predicted base-recognition pocket of *Pf*PucC differs from pyrimidine-producing Pycsar cyclases, providing a structural rationale for GTP selection and cGMP production. This suggests that Pycsar-related cyclases can maintain a conserved catalytic mechanism while undergoing variation of a limited set of residues that determine nucleobase specificity. Such variation appears to have allowed this clade to signal via a range of cyclic mononucleotides, including cGMP, and potentially cAMP.

The *Pf*PucTIR effector revealed an unusual mode of autoinhibition. In the absence of cGMP, *Pf*PucTIR forms a dimer-of-dimers tetrameric assembly in which TIR domains are already positioned close to one another, but remain catalytically inactive. This differs from many TIR-containing immune proteins, where inactivity is often achieved by keeping TIR domains spatially separated until activation promotes self-association^27,28^. In *Pf*PucTIR, inhibition instead appears to arise from self-sequestration in a preassembled but misoriented tetrameric oligomeric state. The TIR domains are proximal, but their relative arrangement does not create an NADase-competent composite active site.

The cGMP-bound structures support a stepwise model for release of this autoinhibited state. The partially occupied cGMP-bound tetramer and dimer appear to capture intermediate-like conformations in which cGMP binding remodels and dissociates the apo assembly but does not yet generate the active filament architecture. Full cGMP occupancy stabilizes an extended domain-swapped dimer that can stack into a linear filament. The structures therefore suggest that *Pf*PucTIR activation is not simply ligand-induced oligomerization, but a staged process in which cGMP binding progressively releases self-sequestration and converts the effector into a filament-forming state. Because these structures represent snapshots, the precise temporal order and relative abundance of these states during infection remain to be established. Nevertheless, they provide a coherent structural pathway linking cGMP binding to effector remodeling and filament assembly. This might also suggest that the proposed progressive assembly could serve a threshold setting mechanism of action of the Pucsar systems.

Filament formation provides a structural rationale for NADase activation. The linear filament is predicted to create composite NAD^+^-binding pockets between TIR domains from adjacent stacked 2:2 cGAMP:PucTIR dimers, while also repositioning the BB loop away from the predicted NAD^+^-binding site. Movement of this loop is a recurring feature of TIR NADase activation, where substrate access depends on an open, catalytically competent TIR interface.^28,29^ Consistent with this model, mutations that disrupt cGMP binding, dimer stabilization, or linear filament contacts strongly reduce NAD^+^ hydrolysis. By contrast, mutations predicted to disrupt the higher-order zig-zag packing do not substantially impair NADase activity *in vitro*. However, these same mutations reduce antiphage defense in cells, indicating that the zig-zag interface is dispensable for NAD⁺ hydrolysis by the purified proteins, but contributes to PucsarTIR function during infection. This distinction suggests that the linear filament contains the catalytically important architecture, whereas the zig-zag arrangement may contribute to filament packing, stability, organization, localization, or productive activation *in vivo* rather than being strictly required for catalysis.

*Pf*PucTIR is phylogenetically related to the Pycsar-associated CNBD-TIR systems and a part of the broader family of assembly-activated TIR NADases. Previous Pycsar studies established that phage infection can stimulate production of cyclic pyrimidine mononucleotides, including cCMP and cUMP, which activate downstream PycTM and PycTIR effectors^12,13^. Structural analysis of PycTIR further showed that cUMP binding to the CNBD promotes oligomerization and supported a model in which ligand-bound PycTIR assemblies extend into filaments^13^. However, the TIR domain was absent from the solved structures of the cUMP-bound complex, leaving the TIR-TIR interfaces and the formation of the NADase-active architecture unresolved. PucsarTIR clarifies this process by capturing a full-length CNBD-TIR effector in multiple states, from an inactive self-sequestered tetramer to partially cGMP-bound intermediate-like assemblies to a fully cGMP-bound filamentous state.

The requirement for higher-order TIR assembly to generate composite NADase active sites is not unique to *Pf*PucTIR. TIR-SAVED, TIR-STING, and SARM1 all illustrate the broader principle that ligand-dependent or activation-dependent TIR oligomerization can reconstitute NADase-active interfaces^26,30–32^ The distinctive feature of *Pf*PucTIR is therefore not filamentation, but the regulatory route by which filamentation is achieved. In *Pf*PucTIR, cGMP does not simply bring separated TIR domains together; instead, it appears to remodel a preassembled, inactive CNBD-TIR tetramer through partially occupied states before stabilizing a fully cGMP-bound dimeric repeat that stacks into the active linear filament. This provides a structural pathway for how a Pucsar-related CNBD-TIR effector can be converted from a self-sequestered inactive state into an NADase-active assembly.

More broadly, *Pf*PucTIR adds to an emerging view that antiphage defense systems often restrain potentially toxic effectors until infection-associated signals trigger their release or activation. This principle is seen, for example, in systems such as Lamassu^33,34^ and PARIS^35^, where nuclease or toxin effectors are held inactive by partner proteins and activated only after detection of phage-associated signals. *Pf*PucTIR achieves a related regulatory outcome through a different strategy. Rather than being sequestered by a separate inhibitory partner, the effector is self-sequestered in an inactive oligomeric state. Binding of cGMP remodels this preassembled complex and promotes stepwise formation of the linear filament required for NADase activity. This provides a layered regulatory mechanism in which cGMP production, ligand occupancy, and filament assembly together help restrict NAD^+^ depletion to infection-associated conditions.

Together, our findings establish Pucsar as a cGMP-producing branch of the Pycsar-related antiphage clade of cNMP cyclases and reveal how this signal is coupled to activation of a TIR NADase effector. The system illustrates how bacteria can adapt a conserved cyclic nucleotide cyclase scaffold to generate a diversity of second messengers, and how this signal can be translated into antiviral activity through conformational remodeling and release of a self-sequestered effector. The structural pathway described here suggests that multiple regulatory layers are built into *Pf*PucTIR activation. The apo effector is inactive despite being oligomeric, partially cGMP-bound states remodel the complex without forming the active architecture, and full cGMP occupancy stabilizes the filament-forming dimeric repeat. This stepwise behavior may help prevent inappropriate activation of a potentially toxic NADase while allowing rapid effector assembly once the cGMP signal is produced. More broadly, the diversity of cNMP cyclases predicted in this work to participate in antiviral immunity raises the possibility that other members of this family may use additional cyclic nucleotide signals and effector mechanisms. The identification of further immune-associated cyclase lineages, including cyclases linked to STAND NTPases, suggests that diversification of cNMP signaling may extend beyond the Pycsar/Pucsar family characterized here. Together, these observations point to a broader repertoire of nucleotide-controlled antiphage defense mechanisms.

## Limitations of the study

The structures described here provide snapshots of distinct *Pf*PucTIR conformational states and support a stepwise model of cGMP-dependent activation. However, the temporal order and relative abundance of these states during phage infection remain to be established. Although structural modelling predicts composite NAD^+^-binding sites within the *Pf*PucTIR filament, we did not directly capture NAD^+^, or the inhibitor BAD. bound to the active assembly, likely reflecting the transient nature of substrate engagement. In addition, while the *Ec*PucsarTM cyclase produces cGMP and the cognate *Ec*PucTM effector is required for antiphage defense, the mechanism by which cGMP regulates *Ec*PucTM and how its hexameric membrane assembly restricts phage propagation remain unresolved. Finally, predicted nucleotide preferences across the broader Pycsar-related cyclase family are based on computational modelling and will require experimental characterization of additional representatives.

## Acknowledgements

We thank Alexander Harms (ETH Zurich) for providing the Basel phage collection. Y.W. is supported by a NIHR Southampton Biomedical Research Centre (BRC) Postdoctoral Bridging Fellowship. D.J.P. is supported by the National Institutes of Health (NIH) grant GM145888, the Maloris Foundation, and the Memorial Sloan Kettering Cancer Center Core Grant (P30-CA008748). We would like to thank M. Jason de la Cruz and Sagnik Sen (MSKCC) for assistance with cryo-EM data collection. In addition to the MSKCC cryo-EM facilities, portions of this work were performed at the National Center for CryoEM Access and Training (NCCAT) and the Simons Electron Microscopy Center at the New York Structural Biology Center. These facilities are supported by the NIH Common Fund Transformative High Resolution Cryo-Electron Microscopy Program (U24 GM129539), the Simons Foundation (SF349247), and the New York State Assembly. This work was also supported by Royal Society grant RGS\R2\222312 to F.L.N. A.M.B. and L.A. are supported by the Intramural Research Program of the National Institutes of Health (NIH). The contributions of the NIH author(s) were made as part of their official duties as NIH federal employees, are in compliance with agency policy requirements, and are considered Works of the United States Government. This work utilized the High Performance Computing Biowulf system. R.H.H is supported by the National Institutes of Health (NIH) grant GM145060.

## Author contributions

Conceptualization, D.J.P., L.A., F.L.N. R.H.H; Methodology, D.J.P., F.L.N., R.H.H, A.C, A.M.B., L.A. S.F; Investigation, A.C., Y.W., A.M.B., Z.Z., S.F.; Visualisation, A.C., Y.W., A.M.B.; Writing – Original Draft, A.C., A.M.B., L.A., F.N., D.P.; Writing – Review & Editing: all authors; Resources, L.A., F.L.N., R.H.H., D.J.P. Funding Acquisition, D.J.P.

## Declaration of interests

The authors declare no competing interests.

## Data and code availability

Cryo-EM maps have been deposited in the Electron Microscopy Data Bank (EMDB) under the accession codes EMDB: EMD-78085 (Apo-*Pf*PucTIR tetramer), EMD-78162 (1:2 cGMP:*Pf*PucTIR dimer), EMD-78166 (2:4 cGMP:*Pf*PucTIR tetramer), EMD-78196 (*Pf*PucTIR filament, focused map), EMD-78189 (*Pf*PucTIR filament, global map) and EMD-78190 (*Ec*PucTM hexamer). The corresponding atomic coordinates of the cryo-EM structures have been deposited in the Protein Data Bank (PDB) under the accession codes PDB: 37DE (Apo-*Pf*PucTIR tetramer), PDB: 37GA (1:2 cGMP:*Pf*PucTIR dimer), PDB: 37GH (2:4 cGMP:*Pf*PucTIR tetramer), PDB: 37HX (2:2 cGMP:*Pf*PucTIR dimer), PDB: 37HQ (*Pf*PucTIR filament) and PDB: 37HR (*Ec*PucTM hexamer). Crystal data of the Apo *Pf*PucC cyclase has been deposited to PDB under the accession code PDB: 37UE.

## Methods

### Sequence analysis

Sequence profile searches to identify cNMP cyclase domains were conducted using PSIBLAST^36^ and hidden Markov model (HMM) searches via JACKHMMER^37^. These queries were performed against either the NCBI non-redundant (nr) protein database^38^ or the nr database clustered at 50% identity (nr50). The MMseqs program^39^ was used to cluster the true positive sequences based on bitscore or percentage similarity. Multiple sequence alignments were generated using either FAMSA^40^ or the MAFFT program (utilizing the local-pair algorithm with parameters: –maxiterate 3000, –op 1.5, and –ep 0.2)^41^.

### Domain identification, comparative genomics and phylogenetic analysis

We identified known domains using HMMs from the Pfam database^42^ alongside a curated collection of in-house HMMs and PSSMs from the Aravind laboratory. PSSMs were searched using RPSBLAST^43^ while HMMs were queried using HMMSCAN^44^. To facilitate domain discovery and remote homology detection, HHpred^45^ was employed against the Pfam^42^ and PDB70^46^ databases. These computational pipelines and the extraction of genomic neighborhoods were automated using the ROTIFER package (http://github.com/leepbioinfo/rotifer). Conserved gene neighborhoods were defined by clustering adjacent genes via the MMseqs program, utilizing a 150-nucleotide maximum distance cutoff and filtering by open reading frame (ORF) orientation. Phylogenetic trees were constructed using FastTree^47^ and iqTREE2^48^.The above-generated alignments were split into subclades guided by the trees obtained in the phylogenetic analysis and used to generate consensus patterns to identify subclade-specific residues.

### Structure Analysis

3D structural modeling was performed using AlphaFold3^21^.We assessed model reliability using the predicted local Distance Difference Test (plDDT) for local chain confidence and the predicted Alignment Error (PAE) to evaluate relative domain orientations. The above-obtained consensus patterns were mapped onto the AF3-generated structures of each subclade and examined with a 4 Angstrom probe to identify protein-nucleotide interactions. For nucleotide-protein interactions, we utilized the atom-by-atom plDDT metric obtained directly from the AF3-generated CIF files to test the difference between purine and pyrimidine NTPs. Tests for significance and analysis of the plDDT scores were performed using the R language. Structure similarity searches were performed with DALI^49^ and Foldseek^50^ and molecular visualization was conducted using MOL*^51^.

### Bacterial strains and phages

*E. coli* strain DH10B was used to clone plasmids pUCP20 with PucsarTIR and pRSFduet with PucsarTM. *E. coli* BL21-AI and *P. aeruginosa* PAO1 were used for *in vivo* tests with PucsarTM and PucsarTIR respectively. Bacterial strains were grown at 37 °C in Lysogeny Broth (LB) with 180 rpm shaking for liquid cultures, or in LB agar (LBA, 1.5% (w/v) agar) plates for solid cultures. Whenever applicable, LB was supplemented with 100 µg/ml of ampicillin or 50 µg/ml of kanamycin (for *E. coli*), or 200 µg/ml of carbenicillin (for *P. aeruginosa*). Phages used in this study are described in **Table S5**.

### Cloning of PucsarTIR and PucsarTM for *in vivo* assays

Plasmids and primers used in this work can be found in **Tables S6** and **S7**, respectively. PucsarTIR from *P. fluorescens* ICMP3636 (168605-170256, LKEI01000069) was cloned into pUCP20 by Gibson assembly. PucsarTM from *E. coli* KTE100 (373743-373749, ASVF01000031) was cloned into pRSF-2 by Gibson assembly. The plasmids were recovered in DH10B cells, extracted using Monarch Plasmid Miniprep Kit (New England Biolabs) and confirmed by sequencing at Macrogen (Netherlands) or Plasmidsaurus (USA). Mutations of Pucsar systems were engineered by around-the-horn PCR or Gibson assembly and confirmed by Sanger sequencing at Macrogen. Plasmids were transformed individually or in combinations into competent BL21-AI cells prepared using the Mix&Go! *E. coli* Transformation Kit (Zymo Research), or electroporated into PAO1 cells prepared as previously described^52^.

### Efficiency of plating assays

Overnight bacterial cultures were mixed with 0.6% top agar and overlaid on LBA plates. For inducible PucsarTIR, bacteria were inoculated in LB containing antibiotics, 1 mM IPTG and 0.2% arabinose and incubated at 37 °C and 180 rpm overnight. Ten-fold serial dilutions of the phage stocks were spotted onto the bacterial lawn and the plates incubated overnight at 37 °C. The phage plaques were counted and used to calculate the efficiency of plating relative to the control. Statistical significance was determined using the multiple comparison function from Two-way ANOVA with a p-value of <0.01.

### Liquid assay

Overnight bacterial cultures were diluted to an optical density at 600 nm (OD_600_) of 0.1 in LB containing antibiotics. The bacterial suspension was distributed into wells of a 96-well plate to which phage dilutions or LB were added. The plates were incubated in a Clariostar Plus plate reader at 37 °C with shaking at 200 rpm, with OD_600_ measured every 5 min for 24 h.

### Intracellular and extracellular NAD⁺ measurement

Overnight cultures were adjusted to an OD_600_ of 0.1. Phages were added at an MOI 10 and 0.1. The mixtures were incubated at 37 °C and 180 rpm and sampled at time point 0, 15, 30 and 60 minutes post infection. For extracellular NAD^+^ measurement, the samples were centrifuged at 12,000 *x* g for 1 minute. For total NAD^+^ measurement, the samples were incubated with chloroform before centrifugation. The supernatants were flash frozen with liquid nitrogen. Promega NAD/NADH-Glo assay kit (G9071) was used to measure the NAD^+^ concentration, and intracellular concentration was determined by subtracting extracellular from total values.

### Cloning, expression, and purification of recombinant *Pf*PucC cyclase, *Ec*PucC cyclase, PucTIR effector and mutant PucTIR proteins

Synthetic genes encoding both PucC cyclase and PucTIR effector proteins from *Pseudomonas fluorescens* (*Pf*), were purchased from IDT. The synthetic genes were cloned into pRSF-1 vector, which carries an N-terminal 6XHis tag followed by a SUMO tag. *E. coli* BL21 (DE3) transformed with expression plasmids were grown at 37 °C until A_600_ reached 0.4−0.6. After cooling to 18 °C, expression was induced with 0.5 mM isopropyl-β-D-thiogalactopyranoside (IPTG), and cells were grown at 18 °C overnight. The cultures were harvested, and the cell pellets were resuspended in lysis buffer (20 mM Tris-HCl pH 8.0, 500 mM NaCl, 5% glycerol). Cells were lysed using sonication (1s pulse ON, 2s pulse OFF), followed by centrifugation at 22,000 RPM for 1 hour at 4 °C to remove cell debris. After filtration using 0.45 μm filter, the supernatant was loaded into HisTrap-FF column (GE Healthcare). The *Pf* proteins (PucC and PucTIR) were eluted using elution buffer (20 mM Tris-HCl, pH 8.0, 500 mM NaCl, 500 mM imidazole, 5% glycerol) and fractions containing SUMO tagged proteins were combined. Following this, SUMO tagged proteins were digested with Ulp overnight and then passed through another HisTrap-FF column to collect flow through that contains untagged proteins. Following this, salt concentration of the flow through was raised to 0.8 M (NH4)2SO4 and loaded onto a HiTrap-Phenyl-HP column (GE Healthcare) with buffer A (20 mM Tris-HCl pH 8.0, 1.25 M (NH_4_)_2_SO4) and buffer B (20 mM Tris-HCl pH 8.0). The eluted proteins were then concentrated and further purified on a Superdex 200 size exclusion chromatography (GE Healthcare) column with storage buffer (20 mM HEPES pH 7.0, 250 mM NaCl, 2% glycerol). Protein purity was assessed by SDS-PAGE with Coomassie staining and samples were flash-frozen in liquid nitrogen and stored at −80 °C. All mutants of PucTIR were purified using the same steps as before except Superose 6 10/200 gl size exclusion column was used for the final step.

### Cloning, expression, and purification of recombinant *Ec*PucTM effector

Synthetic genes encoding PucTM effector protein from *Escherichia coli (Ec)*, were purchased from IDT. The synthetic genes were cloned into pRSF-1 vector, which carries an N-terminal 6XHis tag. *E. coli* C41 (DE3) transformed with expression plasmids were grown at 37 °C until A_600_ reached 0.4−0.6. After cooling to 16 °C, expression was induced with 0.5 mM isopropyl-β-D-thiogalactopyranoside (IPTG), and cells were grown at 16 °C overnight. The cultures were harvested, and the cell pellets were resuspended in lysis buffer (20 mM Tris-HCl pH 8.0, 300 mM NaCl, 5% glycerol). Cells were lysed using a homogenizer. 1% LMNG and 0.1% CHS was added to cell lysate and rotated at 4 °C for 3 hours. The lysate was then centrifuged at 22,000 RPM for 1 hour at 4 °C to remove cell debris. Supernatant was loaded onto Ni-beads and purified by gravity flow using 20 mM Tris-HCl, pH 8.0, 300 mM NaCl, 5% glycerol containing buffer containing 60 mM, 100 mM, 250 mM, 300 mM and 500 mM Imidazole respectively. The best fraction was then passed through a gradient elution using AKTA-FPLC (Cytiva) to further remove impurities. The best fractions were pooled and concentrated followed by passing it through a Superose 6 increase 10/300 gl size exclusion column (Cytiva) using 150 mM NaCl, 25 mM Tris pH 7.5 and a detergent mixture with 0.00033% GDN and 0.001% LMNG/0.0001%CHS.

### *In vitro* reconstitution of *Pf*PucC cyclase

For *in vitro* reconstitution assays, 150 μl reactions with 200 μM each of ATP, GTP, CTP and UTP and 20 μM of recombinant PucC were carried out in 1X PucC-Reaction buffer (50 mM CAPSO, pH 7.4, 50 mM KCl, 5 mM MgCl2 or MnCl2, 1 mM DTT). Reactions were incubated for 1 hour at room temperature. The terminal phosphates were removed using 10 U of Calf Intestinal Phosphatase after the reaction was complete and the reaction mixture was filtered using 10 kDa-cutoff centrifuge filters. The flow-through was analyzed using UPLC and LC-MS.

### *In vitro* NAD^+^ hydrolysis activity of *Pf*PucTIR effector

Commercially purchased cyclic mononucleotides, 3′,5′-cCMP (5 μM), 3′,5′-cGMP (0.5 μM) 3′,5′-cAMP (5 μM) and 3′,5′-cUMP (5 μM) were incubated with 200 μM NAD^+^ and 2 μM PucTIR at 25 °C for 1 hour. Following this, the reaction samples were filtered using 10 kDa centrifuge filters and the flow through containing small molecules was analyzed using UPLC-MS. m/z peaks corresponding to NAD^+^ (substrate), ADP ribose and nicotinamide (product) peaks were analyzed.

### Fluorescence plate reader analysis of NADase activity

NADase activity of purified PucTIR was monitored using a plate reader. 30 μL reactions were aliquoted into 96-well plates. The reactions consisted of buffer (20 mM HEPES-KOH pH 7.5, 100 mM KCl), PucTIR, a range of 1 nM to 10 μM synthetic cyclic nucleotide ligands, and 500 μM of the fluorescent NAD^+^ analog ε-NAD^+^. Fluorescent measurements (300 nm excitation/410 nm emission) of each well were taken after 60 min in a Synergy H1 plate reader (BioTek).

### UPLC analysis of products

For PucC-based assays, UPLC analyses were carried out on a Waters prep LC-MS system with a reversed phase Luna C18 column (250 × 2.0 mm; Phenomenex) with a flow rate of 0.3 ml/min. The column was equilibrated with 100% solvent A (5 mM ammonium acetate, pH 5.3), and the following gradient was applied with solvent B (100% acetonitrile): 0 min, 0% B; 0.5 min, 0% B; 3 min, 0% B; 8.5 min, 5% B; 12 min, 12% B; 20 min, 30% B; 35 min, 50% B; 40 min, 100% B. For PucTIR-based assays, Waters HSS T3, 1.8 μM, 2.1×100 mm column was used with 0.05% TFA as buffer A and acetonitrile+0.04%TFA as buffer B. The total run time was 8 min with a gradient between 1 to 5 min.

### Crystallization and structure determination of *Pf*PucC cyclase

PucC was crystallized at 18 °C using the hanging drop vapor diffusion method. Concentrated protein stocks were diluted in Dilution buffer (20 mM HEPES, pH 7.0, 200 mM NaCl, 2% glycerol) to final concentrations. In all cases, optimized crystals were obtained using 48-well hanging drop tray in 2 μl drops mixed 1:1 over a 200 ul reservoir solution. Crystals of native or selenomethionine substituted PucC grew at 5 mg/ml in 10−12% PEG6000, 100 mM Tris-HCl, pH 7.5, 500 mM NaCl. Crystals appeared after 3-5 days and were cryoprotected using reservoir solution supplemented with 25% glycerol. Native and single-wavelength anomalous dispersion (SAD) data were collected using Advanced Photon Source (APS) and processed using the HKL2000 program. Refinement was done using Phenix^53^. C chain out of the four chains of the PucC tetramer was used as a search model to build other three chains using Coot and Phenix ^53,54^.

### Cryo-EM sample preparation of apo*-Pf*PucTIR tetramer, 2:4 cGMP:*Pf*PucTIR complex, 2:2 cGMP:*Pf*PucTIR filament and *Ec* PucTM hexamer

The purified apo*-*PucTIR was concentrated to 1 mg/ml and 4 µl aliquot of the sample was applied to glow-discharged holey gold grids (UltrAuFoil 300 mesh R1.2/1.3). The grids were blotted for 2 s with a wait time of 10s, a force setting of 0 at 4 °C and 100% humidity, then rapidly frozen in liquid ethane using a Vitrobot Mark IV (FEI).

For the 2:4 cGMP:PucTIR sample, 0.5 mM FOM (fluorinated octyl maltoside, Thermo Fischer Scientific, cat# HR2-40633) was added to the 4 mg/ml of PucTIR tetramer purified at a salt concentration of 150 mM NaCl and applied to the same grids as before. The blotting parameters are same as the ones used for apo-PucTIR tetramer.

For the 2:2 cGMP:PucTIR filament sample, cGMP was rapidly added to 0.8 mg/ml of apo-CNBD-TIR tetramer at a salt concentration of 250 mM NaCl and mixed right before it was applied to glow-discharged holey gold grids (UltrAuFoil 300 mesh R1.2/1.3). The grids were blotted for 1 second with a wait time of 2 seconds and the same force setting and humidity parameters as before.

For the PucTM hexamer sample, protein was concentrated to 3 mg/ml and applied to holey gold grids (UltrAuFoil 300 mesh R1.2/1.3), which were glow discharged for 2s. The grids were blotted for 3 seconds with other parameters similar to apo-PucTIR sample.

### Cryo-EM data collection and processing of apo-*Pf*PucTIR tetramer

Cryo-EM data for the apo-PucTIR effector were collected at the National Center for CryoEM Access and Training (NCCAT) using a Titan Krios transmission electron microscope (FEI) operated at 300 kV and equipped with a Gatan K3 direct electron detector. Automated data acquisition was controlled using the Leginon software suite. Movies were recorded in counting mode with a total electron dose of 55.83 e⁻ Å⁻², a defocus range of -0.8 to -2.2 µm, and a calibrated physical pixel size of 1.073 Å.

A total of 5,284 movies were processed using cryoSPARC (v4.2.1)^55^. Patch-based motion correction and patch-based contrast transfer function (CTF) estimation were performed to correct beam-induced motion and determine CTF parameters. Micrographs exhibiting ice contamination, excessive astigmatism, or poor CTF fit were excluded using threshold-based filtering with the *Manually Curate Exposures* job. Representative high-quality 2D class averages were selected to generate templates for subsequent template-based particle picking. Particles were extracted with a binning factor of two (bin4), yielding a total of 1,478,701 particles. Multiple additional rounds of 2D classification were performed to further remove contaminants and poorly aligned particles. Following *ab initio* reconstruction and heterogeneous refinement into four classes, one well-resolved class corresponding to the apo-PucTIR tetramer containing 286,662 particles were selected. For high-resolution refinement, particles were re-extracted at full pixel size (bin1) followed by homogenous and non-uniform refinements. This workflow yielded an overall resolution of 2.78 Å. The structure was analyzed and images were generated using UCSF Chimera^56^.

### Cryo-EM data collection and processing of the 2:4 and 1:2 cGMP:*Pf*PucTIR complexes

Cryo-EM data for the cGMP-bound PucTIR tetramer in 150 mM NaCl was collected at Memorial Sloan Kettering Cancer Center (MSKCC) using a Titan Krios G4 transmission electron microscope (FEI) operating at 300 kV and equipped with a Falcon 4i direct detector, controlled by EPU software. Movies were recorded in super-resolution mode with a total electron dose of 53 e–/Å², a defocus range of -0.8 to -2.2 µm, and pixel size of 0.725 Å.

A total of 9,237 movies were processed using cryoSPARC (v4.2.1)^55^. Patch-based motion correction and patch-based contrast transfer function (CTF) estimation were performed to correct beam-induced motion and determine CTF parameters. Micrographs exhibiting ice contamination, excessive astigmatism, or poor CTF fit were excluded using threshold-based filtering with the *Manually Curate Exposures* job. Representative high-quality 2D class averages were selected to generate templates for subsequent template-based particle picking. Particles were extracted with a binning factor of two (bin4), yielding a total of 1,544,964 particles. Multiple additional rounds of 2D classification were performed to further remove contaminants and poorly aligned particles. Following *ab initio* reconstruction and heterogeneous refinement into three classes, two well-resolved classes corresponding to the tetramer and dimer containing 67,364 particles and 148,045 particles respectively were selected. For high-resolution refinement, particles were re-extracted at full pixel size (bin1) followed by homogenous and non-uniform refinements. This workflow yielded an overall resolution of 2.8 Å for the 2:4 cGMP:PucTIR complexes and 3.1 Å for the 1:2 cGMP:CNBD-TIR complex.

### Cryo-EM data collection and processing of the 2:2 cGMP-*Pf*PucTIR filament

Cryo-EM data for the cGMP-bound PucTIR filament in 250 mM NaCl were collected at the National Center for Cryo-EM Access and Training (NCCAT) using a Titan Krios transmission electron microscope (FEI) operated at 300 kV and equipped with a Gatan K3 direct electron detector. Automated data acquisition was controlled using the Leginon software suite. Movies were recorded in counting mode with a total electron dose of 52.33 e⁻ Å⁻², a defocus range of -0.8 to -2.2 µm, and a calibrated physical pixel size of 1.073 Å.

A total of 6,819 movies were processed using cryoSPARC (v4.2.1)^55^. Patch-based motion correction and patch-based contrast transfer function (CTF) estimation were performed to correct beam-induced motion and determine CTF parameters. Micrographs exhibiting ice contamination, excessive astigmatism, or poor CTF fit were excluded using threshold-based filtering with the *Manually Curate Exposures* job. The remaining high-quality micrographs were subjected to particle picking using the *Manual Picker* to select filamentous particles, followed by two-dimensional (2D) classification. Representative high-quality 2D class averages were selected to generate templates for subsequent template-based particle picking. Particles were extracted with a binning factor of two (bin2), yielding a total of 565,553 particles. Multiple additional rounds of 2D classification were performed to further remove contaminants and poorly aligned particles. Following *ab initio* reconstruction and heterogeneous refinement into four classes, two well-resolved classes containing a combined total of 467,973 particles were selected and merged for further processing. These particles were refined using homogeneous refinement. For high-resolution refinement, particles were re-extracted at full pixel size (bin1). The corresponding bin2 volume was rescaled and transferred to bin1 using the RELION handler implemented in RELION^57^. The particles were then subjected to global and local CTF refinement to correct residual astigmatism and higher-order aberrations, followed by a final round of non-uniform refinement. This workflow yielded a global filament reconstruction at an overall resolution of 3.34 Å based on 443,336 particles, with substantially improved map quality.

To further enhance density at the interfaces between CNB domains, TIR domains, and the cGMP-binding pocket, local refinement was performed. A local mask encompassing a PucTIR dimer was generated in RELION and applied to the same particle set, resulting in a locally refined map at 3.36 Å resolution. The final global and local reconstructions enabled confident atomic model building of the full-length PucTIR assembly within the higher-order filament, revealing well-defined density for bound cGMP molecules.

### Cryo-EM data collection and processing of the *Ec*PucTM hexamer

Cryo-EM data for the cGMP-bound PucTIR tetramer in 150 mM NaCl was collected at Memorial Sloan Kettering Cancer Center (MSKCC) using a Titan Krios G4 transmission electron microscope (FEI) operating at 300 kV and equipped with a Falcon 4i direct detector, controlled by EPU software. Movies were recorded in super-resolution mode with a total electron dose of 53 e–/Å², a defocus range of -0.8 to -2.2 µm, and pixel size of 0.725 Å.

A total of 11,861 movies were processed using cryoSPARC (v4.2.1)^55^. Patch-based motion correction and patch-based contrast transfer function (CTF) estimation were performed to correct beam-induced motion and determine CTF parameters. Micrographs exhibiting ice contamination, excessive astigmatism, or poor CTF fit were excluded using threshold-based filtering with the *Manually Curate Exposures* job. Representative high-quality 2D class averages were selected to generate templates for subsequent template-based particle picking. Particles were extracted with a binning factor of two (bin4), yielding a total of 1,099,687 particles. Several additional rounds of 2D classification were performed to further remove contaminants and poorly aligned particles. Following *ab initio* reconstruction and heterogeneous refinement into two classes, well-resolved classes corresponding to the hexamer containing 246,004 particles were selected. For high-resolution refinement, particles were re-extracted at full pixel size (bin1) followed by homogenous and non-uniform refinements. This workflow yielded an overall resolution of 2.67 Å for the PucTM hexameric complex. Six unassigned densities are located towards the inside of the hexameric pore likely corresponding to detergents. This density lacked defining features for a specific chemical orientation, and therefore, was excluded from the final atomic model.

### Model building and refinement

Initial atomic models were generated using AlphaFold3 predictions^21^ and docked into cryo-EM density maps. Manual model building was performed in Coot^54^ to adjust side chains, loops, and ligand positions. Real-space refinement was carried out in PHENIX^53^ with geometry restraints. Model quality was assessed using standard validation metrics, including Ramachandran statistics and map-to-model correlation.

### For Proteinix models

Proteinix software^24^ was used to generate 6:6:6 cGMP:NAD^+^:*Pf*PucTIR filament complex (pTM = 0.64, ipTM = 0.61). This was also used to generate 2:2:4 *Bc*PycC:UTP:Mn^2+^ (pTM = 0.96, ipTM = 0.95) and 2:2:4 *Ec*PycC:CTP:Mn^2+^ (pTM = 0.89, ipTM = 0.92).

## Notes

### Competing Interest Statement

The authors have declared no competing interest.

