## Supplementary Figures S1-S7 and Supplementary Tables S2 and S5-S7 for "Cyclase-generated 3’,5’-cGMP activates PucsarTIR antiphage defense through a stepwise zig-zag aligned filament assembly"

### Supplementary information

**Figure S1.** Biochemical characterization of *Pf*PucC and *Ec*PucC protein and analysis of Pucsar antiphage systems.

**Figure S2.** Structural basis for nucleotide recognition by Pycsar and Pucsar cyclases.

**Figure S3.** Cryo-EM structure and oligomeric interfaces of apo-*Pf*PucTIR.

**Figure S4.** Cryo-EM structural analysis of 1:2 and 2:4 cGMP:*Pf*PucTIR complexes.

**Figure S5.** Cryo-EM structure of the cGMP-induced *Pf*PucTIR filament.

**Figure S6.** Mutational analysis of the cGMP-induced *Pf*PucTIR filament.

**Figure S7.** Purification and cryo-EM analysis of the EcPucTM hexamer.

**Table S1**. Cyclase proteins included in the phylogenetic analysis. (*provided separately*)

**Table S2.** AlphaFold 3 analysis of predicted nucleotide preferences across cNMP cyclase subclades.

**Table S3.** Crystallography data collection and refinement statistics. *(provided separately)*

**Table S4.** Cryo-EM data collection and refinement statistics. (*provided separately*)

**Table S5** List of bacterial strains and phages used in this study.

**Table S6** List of plasmids used in this study.

**Table S7** List of primers used in this study.

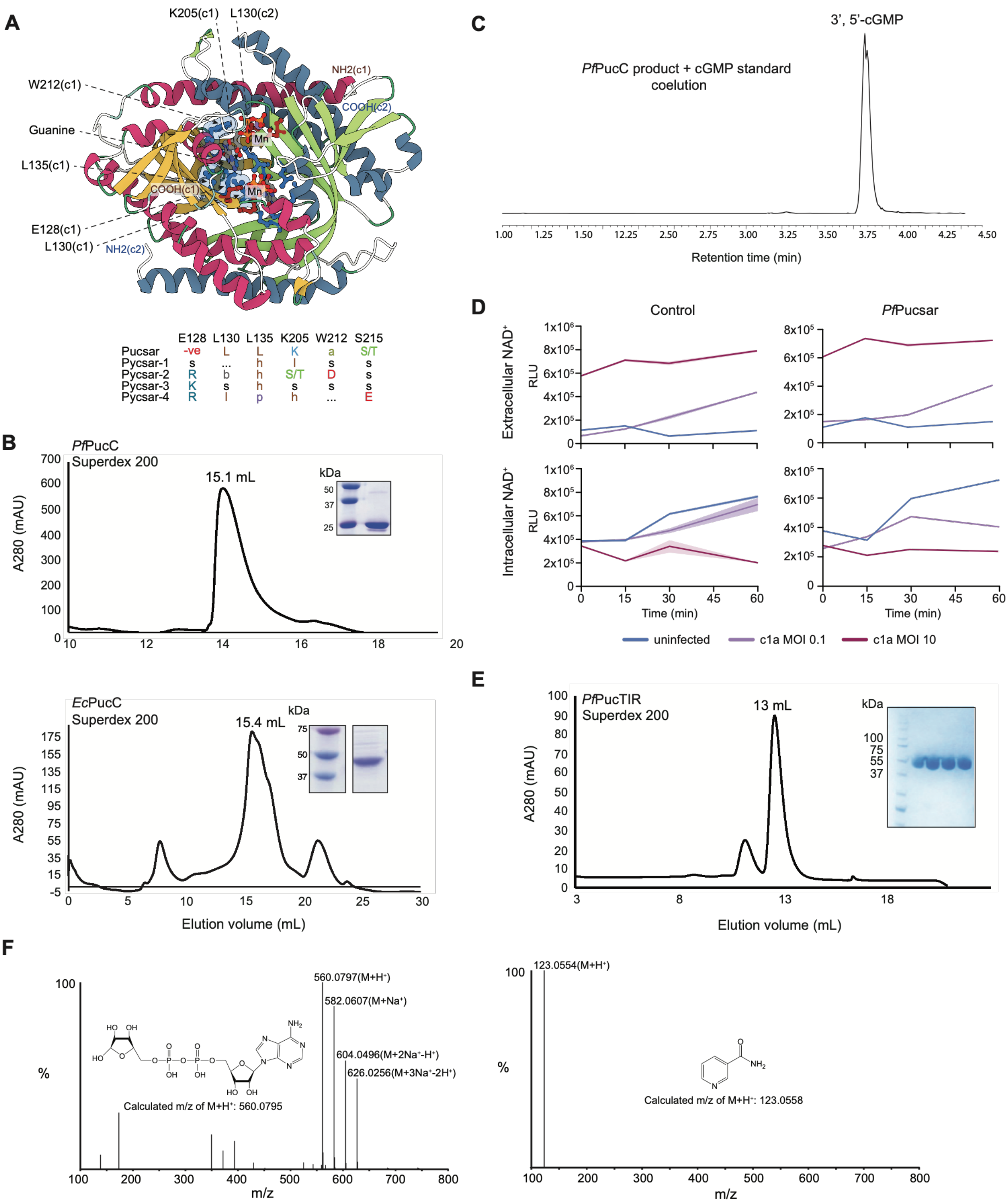


**Figure S1. Biochemical characterization of *Pf*PucC and *Ec*PucC protein and analysis of Pucsar antiphage systems.**

**(A)** Predicted nucleobase-recognition determinants in Pucsar and related Pycsar cyclases. Top, AlphaFold 3 model of the *Yersinia mollaretii* Pucsar cyclase bound to GTP, shown in ball-and-stick representation, and two $Mn^{2+}$ ions, shown in violet. The secondary-structure elements of the two chains are colored differently. The catalytic residues Asp80 and Asp132 are shown in dark red, whereas residues predicted to interact with the guanine base are shown in blue and in ball-and-stick representation. The corresponding residues from chain 1 are additionally shown as a molecular surface to illustrate the base-binding pocket. Labels c1 and c2 indicate chains 1 and 2, respectively. Ser215, which also contributes to the nucleobase-binding pocket, is not shown because of its small side chain. Bottom, residues conserved at the corresponding nucleobase-recognition positions at 90% consensus across the five Pycsar-related subclades, including Pucsar. Absolutely conserved positions are indicated by the one-letter amino-acid code. Other consensus categories are indicated as follows: a, aromatic; l, aliphatic; h, hydrophobic; s, small side chain; b, bulky side chain; p, polar; −ve, negatively charged; and …, no conservation.

**(B)** Size-exclusion chromatography profile of purified *Pf*PucC (left panel) and *Ec*PucC (right panel) on a Superdex 200 column. The volume of the main peaks is indicated. The inserts show the corresponding Coomassie-stained SDS-PAGE analysis.

**(C)** Identification of cGMP produced by the purified *Pf*PucC cyclase by comparison with a synthetic cGMP standard using UPLC-MS.

**(D)** Extracellular and intracellular $NAD^+$ levels in control cells and cells expressing *Pf*Pucsar following infection with phage c1a at multiplicities of infection (MOIs) of 0.1 or 10. Uninfected cultures were analyzed in parallel. $NAD^+$ abundance is shown as relative luminescence units (RLU) over 60 min after infection.

**(E)** Size-exclusion chromatography profile of purified *Pf*PucTIR on a Superdex 200 column. The main peak volume is indicated and the inset shows the corresponding Coomassie-stained SDS-PAGE analysis.

**(F)** Representative mass spectra of the products generated following incubation of *Pf*PucTIR with $NAD^+$. Peaks corresponding to ADP-ribose, including protonated and sodium-adduct species, are shown in the upper spectrum, whereas the lower spectrum shows the nicotinamide product. The calculated mass-to-charge ratios are indicated.

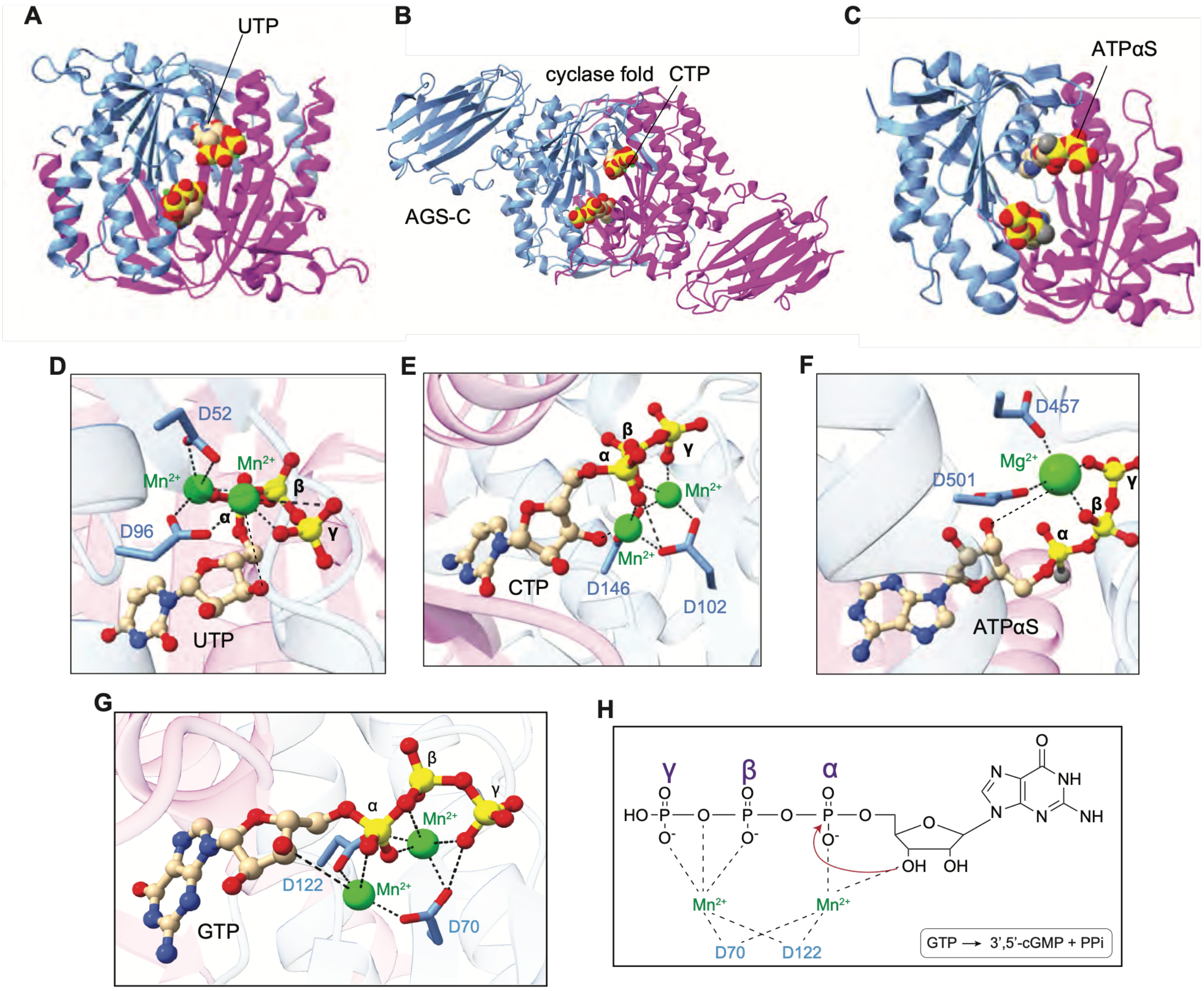


**Figure S2. Structural basis for nucleotide recognition by Pycsar and Pucsar cyclases.**

**(A)** Protenix predicted dimeric complex of *Bc*PycC bound to UTP and $Mn^{2+}$ with a 2:2:4 stoichiometry. The two protein chains are shown in light blue and magenta.

**(B)** Protenix predicted dimeric complex of *Ec*PycC bound to CTP and $Mn^{2+}$ with a 2:2:4 stoichiometry.

**(C)** Crystal structure of the ATP-specific *Ca*RhGC E497K/C566D cyclase variant bound to ATPαS (PDB: 5OYH).

**(D-F)** Enlarged views of the catalytic sites of *Bc*PycC (panel D), *Ec*PycC (panel E), and *Ca*RhGC E497K/C566D (panel F), highlighting the conserved acidic residues involved in divalent-metal coordination and the positions of the α-, β- and γ-phosphates.

**(G)** GTP positioned in its binding pocket in the AF3 model of GTP bound to *Pf*PucC.

**(H)** Schematic summarizing the coordination of D170 and D122 with metals, phosphate and 3’-OH of GTP in the AF3 model of GTP bound to *Pf*PucC that facilitates attack of the 3’-OH on the $\alpha$-phosphate of GTP to produce cGMP.

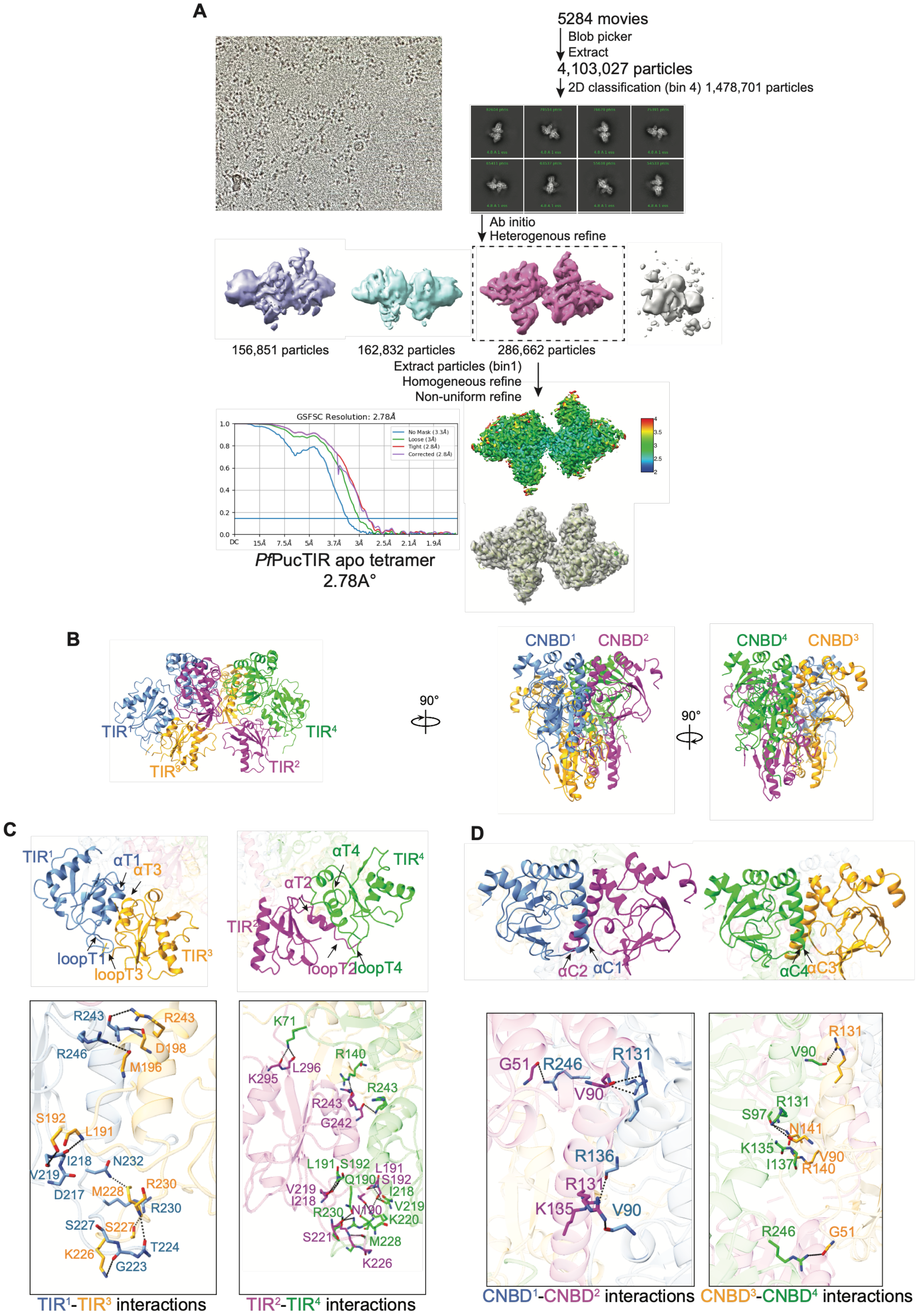
A
5284 movies
Blob picker
Extract
4,103,027 particles
2D classification (bin 4) 1,478,701 particles
Ab initio
Heterogenous refine
156,851 particles
162,832 particles
286,662 particles
Extract particles (bin1)
Homogeneous refine
Non-uniform refine
GSFSC Resolution: 2.78Å
No Mask (3.3Å)
Loose (3Å)
Tight (2.8Å)
Corrected (2.8Å)
PfPucTIR apo tetramer
2.78A°
B
TIR¹
TIR³
TIR²
TIR⁴
90°
CNBD¹
CNBD²
CNBD⁴
CNBD³
C
αT1
αT3
loopT1
loopT3
αT4
αT2
loopT2
loopT4
D
αC2
αC1
αC4
αC3
TIR¹-TIR³ interactions
TIR²-TIR⁴ interactions
CNBD¹-CNBD² interactions
CNBD³-CNBD⁴ interactions

**Figure S3. Cryo-EM structure and oligomeric interfaces of apo-*Pf*PucTIR.**

**(A)** Cryo-EM data processing workflow for apo-*Pf*PucTIR, producing a tetrameric reconstruction at 2.78 Å resolution. Representative micrograph, 2D class averages, intermediate reconstructions, gold-standard Fourier shell correlation (GS-FSC) curve, local-resolution map and fitted atomic model are shown.

**(B)** Structure of the apo-*Pf*PucTIR tetramer, with the four TIR domains coloured blue, magenta, orange and green.

**(C)** Expanded views show the $TIR^1$-$TIR^3$ and $TIR^2$-$TIR^4$ interfaces, including the α-helices and loop regions contributing to tetramer formation. Selected interfacial residues and polar contacts are shown in the lower panels.

**(D)** Expanded views show the $CNBD^1$-$CNBD^2$ and $CNBD^3$-$CNBD^4$ interfaces and the corresponding α-helices. For both panels C and D, expanded segments boxed below show selected residues contributing to each interface, while dashed lines indicate predicted polar interactions, and superscript numbers identify the individual protomers.

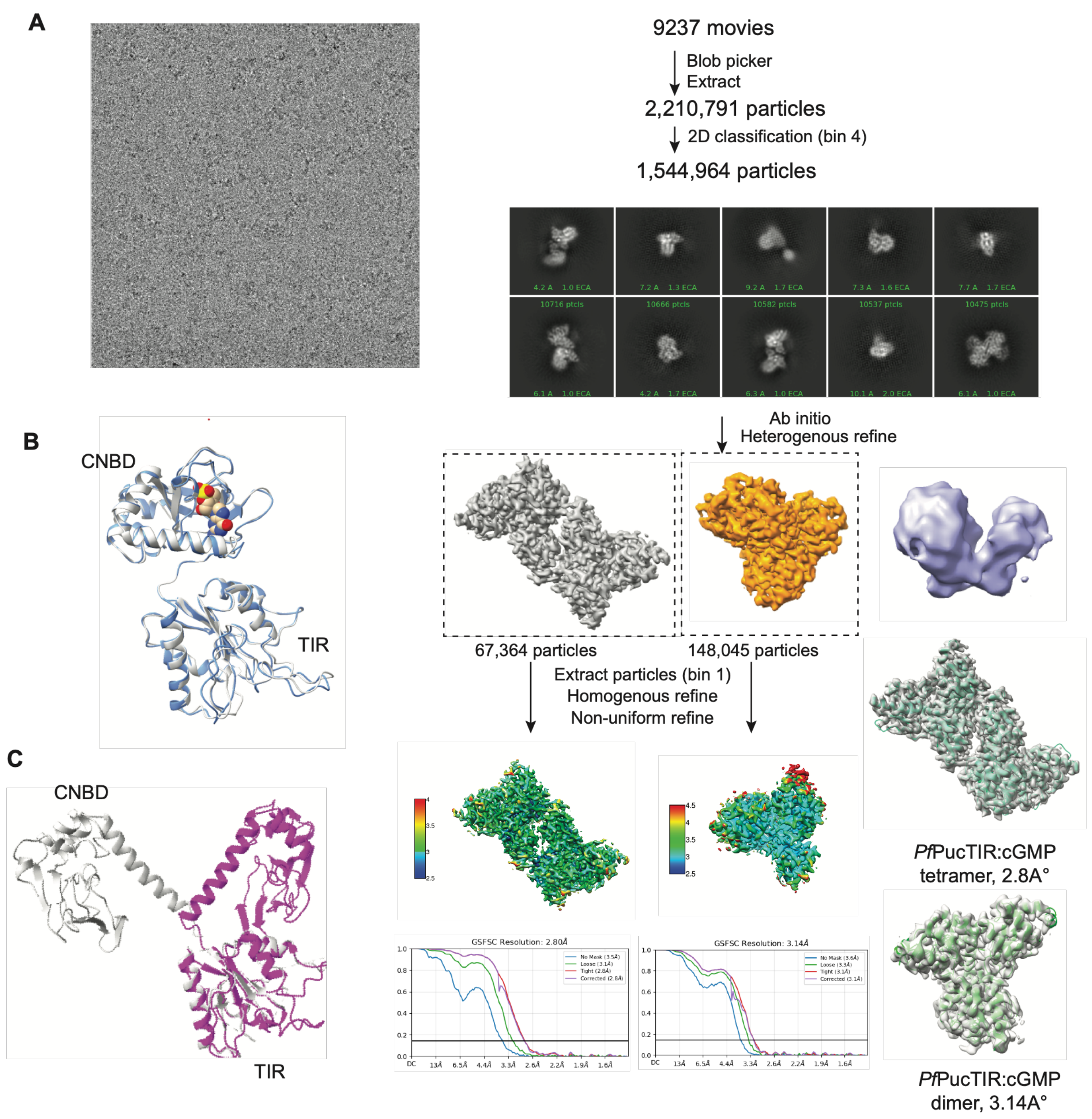


**Figure S4. Cryo-EM structural analysis of 1:2 and 2:4 cGMP:*Pf*PucTIR complexes.**

**(A)** Cryo-EM data processing workflow for cGMP-bound *Pf*PucTIR, producing maps of 2:4 cGMP:PucTIR at 2.80 resolution and 1:2 cGMP:PucTIR at 3.14 Å resolution. Representative micrograph, 2D class averages, reconstructions, GS-FSC curves, local-resolution maps and fitted atomic models are shown.

**(B)** Superposition of the fold-back protomer from apo-*Pf*PucTIR (in silver) with the corresponding fold-back protomer from the 1:2 cGMP:PucTIR dimer (in blue), aligned on the TIR domains, exhibiting a small rmsd = 1.02 Å.

**(C)** Superposition of the extended protomer from apo-*Pf*PucTIR (in silver) with the corresponding protomer from the 1:2 cGMP:PucTIR dimer, showing a larger conformational rearrangement associated with partial cGMP binding (rmsd = 5.32 Å).

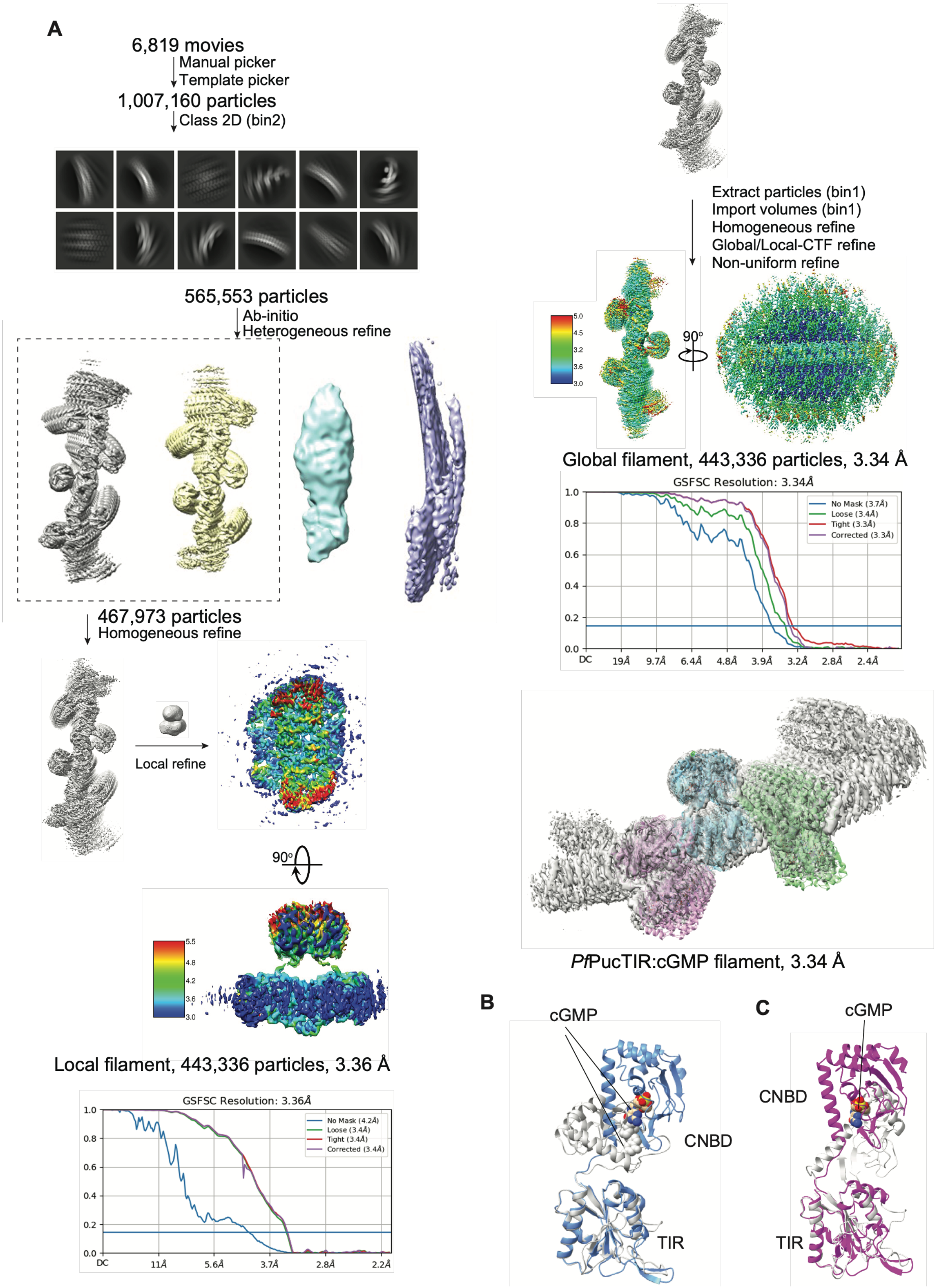

**Figure S5. Cryo-EM structure of the cGMP-induced *Pf*PucTIR filament.**

**(A)** Cryo-EM data processing workflow for cGMP-bound *Pf*PucTIR filaments, producing a map at 3.34 Å resolution. Representative micrograph, 2D class averages, intermediate reconstructions, local resolution maps, gold-standard Fourier shell correlation curves and the fitted atomic model are shown.

**(B)** Superposition of the fold-back protomer from the partially occupied 1:2 cGMP:PucTIR dimer (in silver) and the corresponding protomer from the fully occupied filament-forming 2:2 cGMP-PucTIR dimer (in blue) exhibit an rmsd = 5.6 Å.

**(C)** Superposition of the extended protomer from the partially occupied 1:2 cGMP:PucTIR dimer (in silver) and the corresponding protomer from the fully occupied filament-forming 2:2 cGMP-PucTIR dimer (in magenta) exhibit an rmsd = 5.7 Å.

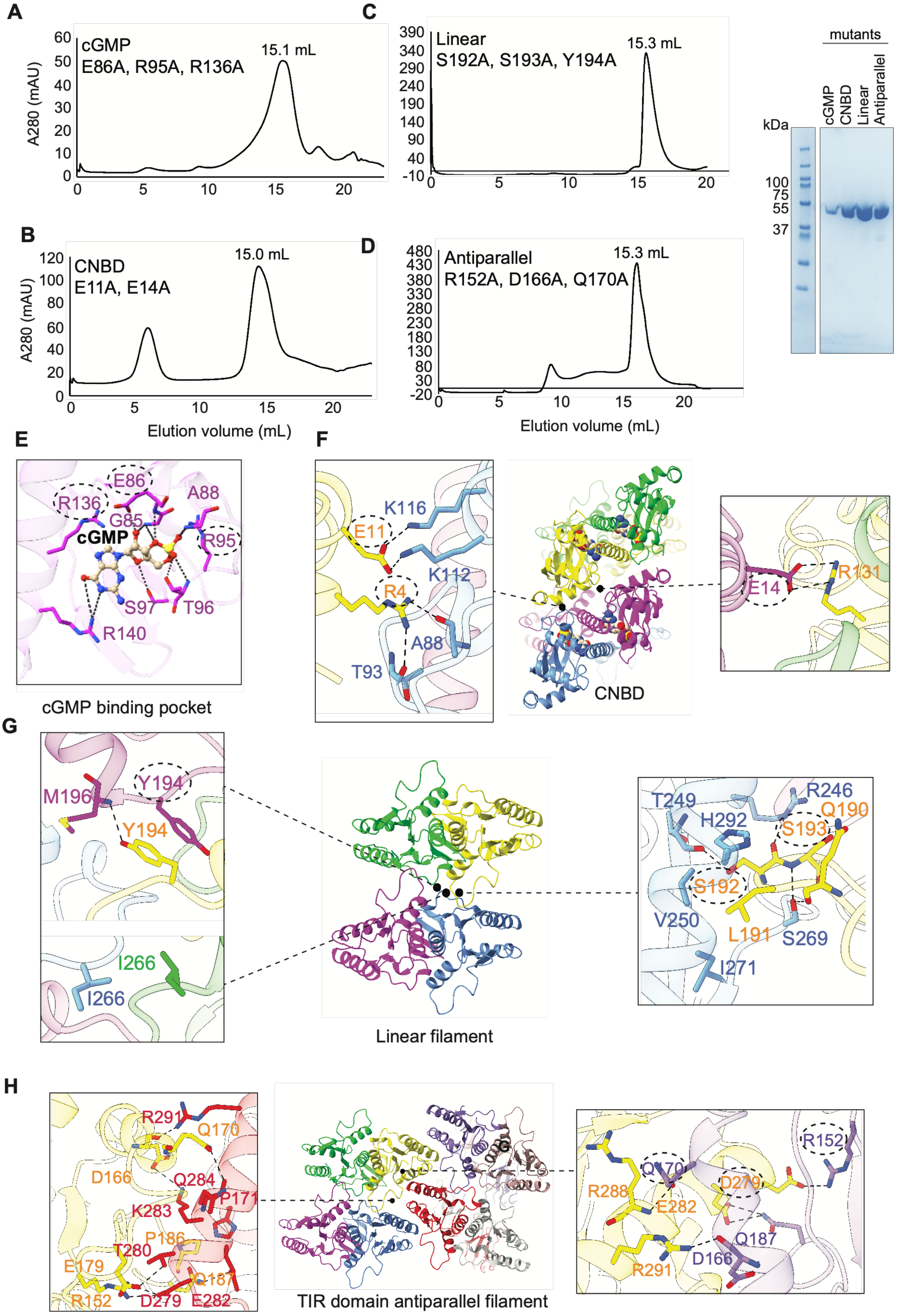

A
cGMP
E86A, R95A, R136A
15.1 mL
A280 (mAU)
C
Linear
S192A, S193A, Y194A
15.3 mL
mutants
cGMP
CNBD
Linear
Antiparallel
kDa
100
75
55
37
B
CNBD
E11A, E14A
15.0 mL
D
Antiparallel
R152A, D166A, Q170A
15.3 mL
Elution volume (mL)
E
F
cGMP binding pocket
CNBD
G
Linear filament
H
TIR domain antiparallel filament

**Figure S6. Mutational analysis of the cGMP-induced *Pf*PucTIR filament.**

**(A-D)** Size-exclusion chromatography profiles and Coomassie-stained SDS-PAGE of purified triple mutants targeting the cGMP-binding pocket (E86A/R95A/R136A) (panel A), the CNBD interface (E11A/E14A) (panel B), the linear filament interface (S192A/S193A/Y194A) (panel C), and the antiparallel filament interface (R152A/D166A/Q170A) (panel D). The principal elution volumes are indicated.

**(E-H)** Close-up view of the cGMP-binding pocket, showing residues involved in nucleotide recognition (panel E); CNBD-CNBD contacts involving Glu11 and Glu14 within the filament assembly (panel F); Interactions contributing to the linear filament interface, including contacts mediated by Ser192, Ser193, and Tyr194 (panel G); Interactions between TIR domains and the antiparallel filament interface, highlighting Arg152, Asp166, and Gln170 (panel H).

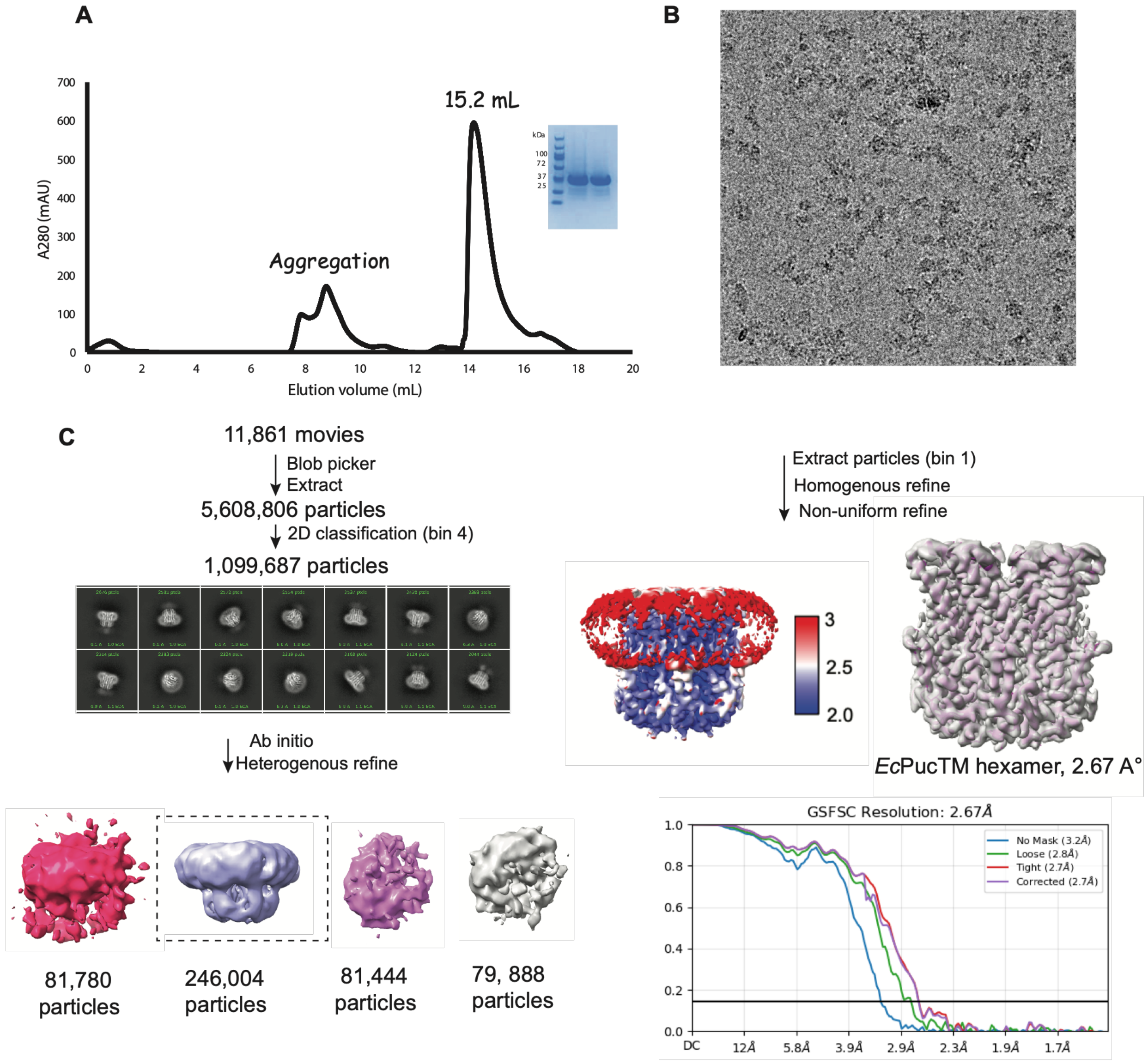


**Figure S7. Purification and cryo-EM analysis of the EcPucTM hexamer.**

**(A)** Size-exclusion chromatography profile of purified *Ec*PucTM in detergent. The major elution peak corresponding to the purified *Ec*PucTM complex is indicated; the inset shows the corresponding Coomassie-stained SDS-PAGE analysis.

**(B,C)** Cryo-EM data-processing workflow for apo-*Ec*PucTM, including a representative micrograph, selected 2D class averages, particle classification and refinement steps leading to the final hexameric reconstruction at 2.67 Å resolution. The final reconstruction, local-resolution map and gold-standard Fourier shell correlation (GS-FSC) curve are shown.

**Table S2.** AlphaFold 3 analysis of predicted nucleotide preferences across cNMP cyclase subclades.

| | Pucsar | Pycsar-1 | Pycsar-2 | Pycsar-3 | Pycsar-like-4 |
|---|---|---|---|---|---|
| **Ions/subunit** | 2 $Mn^{2+}$ | 2 $Mn^{2+}$ | 2 $Mn^{2+}$ | 2 $Mn^{2+}$ | 1 $Mn^{2+}$, 1 $Ca^{2+}$ |
| **ATP preferred** | 0 | 0 | 0 | 0 | 52 |
| **GTP preferred** | 30 | 41 | 5 | 6 | 0 |
| **UTP preferred** | 5 | 97 | 84 | 26 | 4 |
| **Undecided** | 6 | 24 | 21 | 2 | 2 |
| **Total sequences** | 41 | 162 | 110 | 34 | 58 |
| **Binomial test p-value[1]** | 1.12E-05 | 1.04E-06 | 2.20E-16 | 2.67E-04 | 5.50E-12 |
| **Mean ΔpLDDT[2]** | 1.773 | -1.634 | -2.164 | -2.704 | 7.592 |
| **95% CI low** | 0.717 | -2.204 | -2.871 | -4.096 | 6.262 |
| **95% CI high** | 2.829 | -1.063 | -1.458 | -1.312 | 8.922 |
| **ΔpLDDT t-test p-value[3]** | 0.000783 | 3.45E-08 | 9.41E-09 | 0.000192 | 1.11E-16 |

[1] Test of significance of the directionality of called ligands (GTP/ATP vs. UTP).

[2] Structure-level mean of changes in ligand atom pLDDT scores across the two conditions (mean of GTP/ATP - mean of UTP).

[3] One-sided, one-sample t-test evaluating significance of pLDDT mean score difference from zero in the expected direction.

**Table S5** List of bacterial strains and phages used in this study.

| Name | Reference | Sequence |
|---|---|---|
| *Strain* | | |
| *E. coli* DH10B | NEB (cat. C3019) | NZ_CP110018.1 |
| *E. coli* BL21-AI | Thermo Fisher (cat. C607003) | NZ_CP047231.1 |
| *P. aeruginosa* PAO1 | Fagenbank | NC_002516.1 |
| *Phage* | | |
| *Escherichia* phage Lambda-vir | Fagenbank | NC_001416 [a] |
| *Escherichia* phage T1 | Fagenbank | NC_005833.1 |
| *Escherichia* phage T2 | DSMZ | NC_054931.1 |
| *Escherichia* phage T3 | Fagenbank | NC_003298.1 |
| *Escherichia* phage T4 | Fagenbank | NC_000866.4 |
| *Escherichia* phage T5 | Fagenbank | AY543070.1 |
| *Escherichia* phage T6 | Fagenbank | MH550421.1 |
| *Escherichia* phage T7 | Fagenbank | NC_001604.1 |
| *Bas35* | Alexander Harms | MZ501113 |
| *Bas36* | Alexander Harms | MZ501096 |
| *Bas37* | Alexander Harms | MZ501089 |
| *Bas38* | Alexander Harms | MZ501052 |
| *Bas43* | Alexander Harms | MZ501106 |
| *Bas47* | Alexander Harms | MZ501047 |
| *Bas64* | Alexander Harms | MZ501081 |
| *Bas66* | Alexander Harms | N/A |
| *Bas68* | Alexander Harms | MZ501055 |
| *Bas67* | Alexander Harms | MZ501064 |

**Table S6** List of plasmids used in this study.

| Plasmid | Description | Promoter | Origin of replication | Resistance | Cloning method |
|---|---|---|---|---|---|
| pUOS370 | pUCP20-PucsarTIR | lac | pRO1600 oriV | Carb | Gibson assembly |
| pUOS371 | pRSF-PucsarTM | native | RSF | Kan | Gibson assembly |
| pUOS372 | pUCP20-PucsarTIR S192A/S193A/Y194A | lac | pRO1600 oriV | Carb | Around-the-horn on pUOS370 |
| pUOS373 | pUCP20-PucsarTIR R152A/D166A/Q170A | lac | pRO1600 oriV | Carb | Around-the-horn on pUOS370 |
| pUOS374 | pUCP20-PucsarTIR E86A/R95A/R136A | lac | pRO1600 oriV | Carb | Around-the-horn on pUOS370 |
| pUOS375 | pUCP20-PucsarTIR E11A/E14A | lac | pRO1600 oriV | Carb | Around-the-horn on pUOS370 |
| pRSF-2 | pRSF-PucsarTM | native | RSF | Kan | Gibson assembly |
| pUOS376 | pRSF-PucsarTM Q16A | native | RSF | Kan | Around-the-horn on pUOS371 |
| pUOS377 | pRSF-PucsarTM Y166A | native | RSF | Kan | Around-the-horn on pUOS371 |
| pUOS378 | pRSF-PucsarTM K4A | native | RSF | Kan | Around-the-horn on pUOS371 |
| pUOS379 | pRSF-PucsarTM F98A | native | RSF | Kan | Around-the-horn on pUOS371 |
| pRSF-*Pf*PucC | pRSF-*Pf*Pucsar cyclase | lac | RSF | Kan | Restriction digestion/ligation |
| pRSF-*Ec*PucC | pRSF-*Ec*Pucsar cyclase | lac | RSF | Kan | Restriction digestion/ligation |
| pRSF-*Pf*PucTIR | pRSF-*Pf*PucsarTIR | lac | RSF | Kan | Restriction digestion/ligation |
| pRSF-*Ec*PucTM | pRSF-*Ec*PucsarTM | lac | RSF | Kan | Restriction digestion/ligation |
| pRSF-*Pf*PucTIR1 | pRSF-PucsarTIR S192A/S193A/Y194A | lac | RSF | Kan | Q5 mutagenesis on pRSF-*Pf*PucTIR |
| pRSF-*Pf*PucTIR2 | pRSF-PucsarTIR R152A/D166A/Q170A | lac | RSF | Kan | Q5 mutagenesis on pRSF-*Pf*PucTIR |
| pRSF-*Pf*PucTIR3 | pRSF-PucsarTIR E86A/R95A/R136A | lac | RSF | Kan | Q5 mutagenesis on pRSF-*Pf*PucTIR |
| pRSF-*Pf*PucTIR4 | pRSF-PucsarTIR E11A/E14A | lac | RSF | Kan | Q5 mutagenesis on pRSF-*Pf*PucTIR |

**Table S7** List of primers used in this study.

| Name | Description | Sequence (5'--3') |
|---|---|---|
| FN2287<br>FN2288 | Amplify PucsarTIR from *P. fluorescens* ICMP3636 | acgttgtaaaacgacggccagtgcccaaaaatctcattgaacctataattttca<br>agtcgacctgcaggcatgcaagcttttaggacttacggatttcaaatccc |
| FN2289<br>FN2290 | Amplify pUCP20 backbone | aagcttgcatgcctgcagg<br>ggcactggccgtcgtttt |
| FN1878<br>FN1879 | Amplify PucsarTM from *Escherichia coli* KTE100 | cccaaggggttatgctagttattgcttaacttttacccattagatatccag<br>ttatgcgactcctgcattaggaaattatttcccatcataaaatagttgattgt |
| FN1880<br>FN1752 | Amplify pRSF backbone | atttcctaatgcaggagtcgc<br>gcaataactagcataacccc |
| FN2579<br>FN2580 | Mutate TIR S192A/S193A/Y194A on pUCP20-PucsarTIR | cggctgctagttggaatatctgcggcatgctccacggggc<br>attccaactagcagccgcccccatggatgaccttgaagcggagctgacc aactca |
| FN2581<br>FN2582<br>FN2583 | Mutate TIR R152A/D166A/Q170 A on pUCP21-PucsarTIR | agatcgcaaagatggcatgc<br>atcgcgatttcgctcatg<br>agtcatgagcgaaatgcgcgattccgctggcgacgattgagagcgcttcg<br>gcagaagagatcgcaaagatggcatgctt |
| FN2575<br>FN2576<br>FN2577<br>FN2578 | Mutate TIR E86A/R95A/R136A on pUCP22-PucsarTIR | tagtggctgtcgtgacgtggacggcagcaatcgcaccgacgtgattgccagcttcgaaattc<br>atctcgcaaaagcgatcgccgtgcgcaacgccaacgtcgcccgaccg<br>cgatcgcttttgcgagatcccttgcgaaggccagcga<br>acgtcacgacagccactagtaccgtgatcgctcgcgaa |
| FN2573<br>FN2574 | Mutate TIR E11A/E14A on pUCP23-PucsarTIR | atgagctgtgcgaccagcgcgtcccggtctttccttaatcgctccagcatag<br>gctggtcgcacagctcatggtgcaaccgctggtcaaaca |
| FN2584<br>FN2585 | Mutate TM Q16A on pRSF-duet-PucsarTM | tgtactcgcaacatcactgatagattgatagagaaattctattttttgtcc<br>gtgatgttgcgagtacaattagggcgattgatgttaaaattggtttt |
| FN2592<br>FN2593 | Mutate TM Y166A on pRSF-duet-PucsarTM | ggatagccagacagcaataaacttcgtgcaaatagcgtatcggcagaa<br>attgctgtctggctatccttaagtattggattgtatctttactccttaactatc |
| FN2590<br>FN2591 | Mutate TM K4A on pRSF-duet-PucsarTM | attctattgctttgtccattttgatcactataaaaaataatatacactcga<br>ggacaaagcaatagaatttctctatcaatctatcagtgatgttcagagtacaat |
| FN2586<br>FN2587 | Mutate TM F98A on pRSF-duet-PucsarTM | caccgtaagctattcctttcggtttctgcccataaatcccagaagtag<br>aaggaatagcttacggtgatggagtttttaaagttgagtattcatttattgga |
| PucTIR1<br>PucTIR2 | Mutate TIR S192A/S193A/Y194A on pRSF-Duet-PucsarTIR | gagcatgccgcagatttttcagctggcggcggcgccgatggatgatctggaag<br>cttccagatcatccatcggcgccgccgccagctgaaaaatctgcggcatgctc |
| PucTIR3<br>PucTIR4 | Mutate TIR R152A/D166A/Q170A on pRSF-Duet-PucsarTIR | gcgcgaaaaacatgcgatttttgcgattagcagcgcggaagcgctgagcattgtggc gagcggcattgcgcattttgcgcatgatagcg<br>cgctatcatgcgcaaaatgcgcaatgccgctcgccacaatgctcagcgcttccg cgctgctaatcgcaaaaatcgcatgtttttcgcgc |
| PucTIR5<br>PucTIR6 | Mutate TIR E86A/R95A/R136A on pRSF-Duet-PucsarTIR | gcgcgaaaaacatgcgatttttgcgattagcagcgcggaagcgctgagcattgtggc gagcggcattgcgcattttgcgcatgatagcg<br>cgctatcatgcgcaaaatgcgcaatgccgctcgccacaatgctcagcgcttccg cgctgctaatcgcaaaaatcgcatgtttttcgcgc |

| | | |
|---|---|---|
| PucTIR7<br>PucTIR8 | Mutate TIR E86A/R95A/R136A on pRSF-Duet-PucsarTIR | tttgcgcgcgatctggcgaaagcgattgcggtgcgcaacgcgaac<br>gttcgcgttgcgcaccgcaatcgctttcgccagatcgcgcgcaaa |
| PucTIR9<br>PucTIR10 | Mutate TIR E11A/E14A on pRSF-Duet-PucsarTIR | ctgcgcaaagatcgcgatgcgctggtggcgcagctgatggtgcagccgctggtgaaacaggatcgc<br>gcgatcctgtttcaccagcggctgcaccatcagctgcgccaccagcgcatcgcgatctttgcgcag |
